# A tabular foundation model rivals established species distribution models from presence-only records

**DOI:** 10.64898/2026.11.06.765003

**Authors:** Hideyuki Doi, Takeshi Osawa, Narumasa Tsutsumida

**Affiliations:** Graduate School of Informatics, Kyoto University, Yoshida-honmachi, Sakyo-ku, Kyoto, 606-8501, Japan; Graduate School of Urban Environmental Sciences, Tokyo Metropolitan University, Minami-Osawa 1-1, Hachioji, Tokyo, 192-0397, Japan; Graduate School of Science & Engineering, Saitama University, 255 Shimo-Okubo, Sakura-ku, Saitama City, Saitama, 338-8570, Japan; Space Data Frontiers Research Center, Fujitsu Research, Fujitsu Limited, 4-1-1 Kamikodanaka, Nakahara-ku, Kawasaki, Kanagawa 211-8588, Japan

**Keywords:** distribution prediction, presence-only data, foundation model, species distribution model, TabPFN

## Abstract

Predicting where something occurs from records of where it has been observed, without reliable records of absence, is a common problem in science. Species distribution models (SDMs) such as MaxEnt address it in fields from biological conservation and climate-change assessment to the mapping of disease vectors and archaeological sites, but must be tuned for each dataset. Tabular foundation models, pre-trained on synthetic prediction tasks, predict new datasets without per-dataset training. Only one study has tested them as SDMs, and it fine-tuned on survey data from the evaluated regions. We evaluated tabular foundation models on the standard SDM benchmark (225 species in six regions, with independent presence–absence surveys), under the protocol used to compare 21 methods, and on 120 virtual species with known distributions. Without tuning, a tabular foundation model, TabPFN 3.5, equalled the best published approach, a five-model ensemble (mean AUC 0.732). Fine-tuned on presence–background tasks from other regions, it became the best single model on the benchmark (0.736), outperformed MaxEnt (0.728) and boosted regression trees after correction for multiple comparisons, and was better calibrated than MaxEnt, although six regions were too few to establish the advantage for new regions. Fine-tuning on presence–absence surveys of other species helped only when they came from the evaluated regions. On virtual species, it recovered simple additive relationships less well than regression-based methods but complex ones at least as well when records were unbiased, and was worse with environmentally biased records. Tabular foundation models are competitive SDMs without per-dataset tuning, but they are sensitive to environmentally biased records.

## Introduction

Many scientific questions require predicting where something occurs when the only reliable data are records of where it has been observed. Museum specimens, citizen-science sightings, disease case locations and archaeological finds all record presences, whereas systematic records of absence are rare. In machine learning this is known as learning from positive and unlabelled data (1–3). In ecology, it is addressed by species distribution models (SDMs), which contrast presence records with background points describing the available environment and predict the relative likelihood of occurrence across a landscape (4–7).

SDMs are among the most widely used quantitative tools in environmental science. They inform conservation planning (8), projections of extinction risk under climate change (9) and assessments of biological invasions (10), and the same models map pathogens and their vectors (11, 12), including dengue and its *Aedes* vectors worldwide (13, 14), marine species (15), past distributions (16), archaeological sites (17) and crop suitability (18). MaxEnt (19), the most popular method, has become a de facto standard. A method that predicts better than MaxEnt without requiring more expertise could therefore benefit many fields at once.

The most influential benchmark of these methods contains presence-only records of 225 species in six regions, with independent presence–absence surveys for evaluation (20, 21). Its re-analysis with 21 current methods found no single best method (22). The best results came from a five-model ensemble, down-sampled random forests, boosted regression trees (BRT) and MaxEnt, each of which depends on choices of regularisation, tree complexity or class weighting that strongly affect performance (23–25). Accuracy was lowest for species with few records (26, 27), often those of greatest conservation concern (28).

Foundation models offer a different route. They are large neural networks pre-trained on broad data and applied to new tasks with little task-specific training (29). TabPFN, a transformer pre-trained on millions of synthetic tabular datasets, predicts by in-context learning: the training table is passed to the network together with the points to be predicted, and predictions are returned in a single forward pass, without fitting parameters or tuning hyperparameters (30, 31). On small tables it matched or exceeded tuned gradient-boosted trees (30), and later versions improve accuracy (32, 33). These properties suit SDMs, whose datasets are small (a median of 57 presence records per species here) and whose users often lack the time or expertise to tune each model. Presence–background data nevertheless differ from the tasks on which TabPFN was trained: the classes are highly imbalanced, the “absences” are background points of unknown status, and presence records carry sampling bias (6, 7).

Whether foundation models can serve as SDMs has received little attention. The only previous study, an unreviewed preprint, reported large gains after fine-tuning TabPFN 2.5 (34), but it compared the model with five methods rather than the full benchmark and fine-tuned on presence–absence surveys of other species from the evaluated regions, whose survey sites are shared within biological groups. No study has evaluated foundation models against all published methods, with no information from the evaluated region and under spatially separated testing, or asked whether they recover true species–environment relationships, which requires virtual species with known distributions (35, 36).

Here we evaluate two versions of TabPFN, 3.5 (33) and 2.5 (32), and another tabular foundation model, TabICL (37), on this benchmark under exactly the published protocol (22), after re-fitting the reference models to confirm that our results reproduce theirs. We ask whether an off-the-shelf foundation model competes with established SDMs; whether fine-tuning on other regions improves it, and which training signal transfers; how fine-tuning on surveys from the evaluated regions changes the apparent gain; whether the conclusions hold under spatially separated evaluation (38); and, using 120 virtual species, whether foundation models recover true probabilities of occurrence and responses to environmental gradients.

## Results

### A foundation model matches established SDMs without tuning

Our re-fitted reference models reproduced the published regional mean AUCs of Valavi et al. (22). Across nine methods and six regions, the mean absolute difference was 0.004 (0.001–0.006 for eight methods and 0.013 for default random forests; Fig. S1). Our results are therefore directly comparable with all 21 published methods.

Used off the shelf with balanced bagging (100 bags, each with the presences and an equal number of background points), TabPFN 3.5 reached a mean AUC of 0.732 across the 225 species. This was the same as the mean of the best published approach, the five-model ensemble (0.732), and numerically higher than the means of all published single methods, the best of which were down-sampled random forests (RF down-sampled, 0.730), MaxEnt (0.728) and BRT (0.725; Fig. S12). The differences from the best single methods were not significant: compared with our re-fitted MaxEnt, for example, the difference was +0.004 (95% interval from a bootstrap of species within regions, −0.001 to 0.009; Shaffer-corrected *P* = 0.98). TabPFN 2.5 (0.727) performed similarly to MaxEnt. TabICL (0.706), a second tabular foundation model, ranked below all re-fitted reference models except default random forests (Fig. 1A).

**Figure 1.**
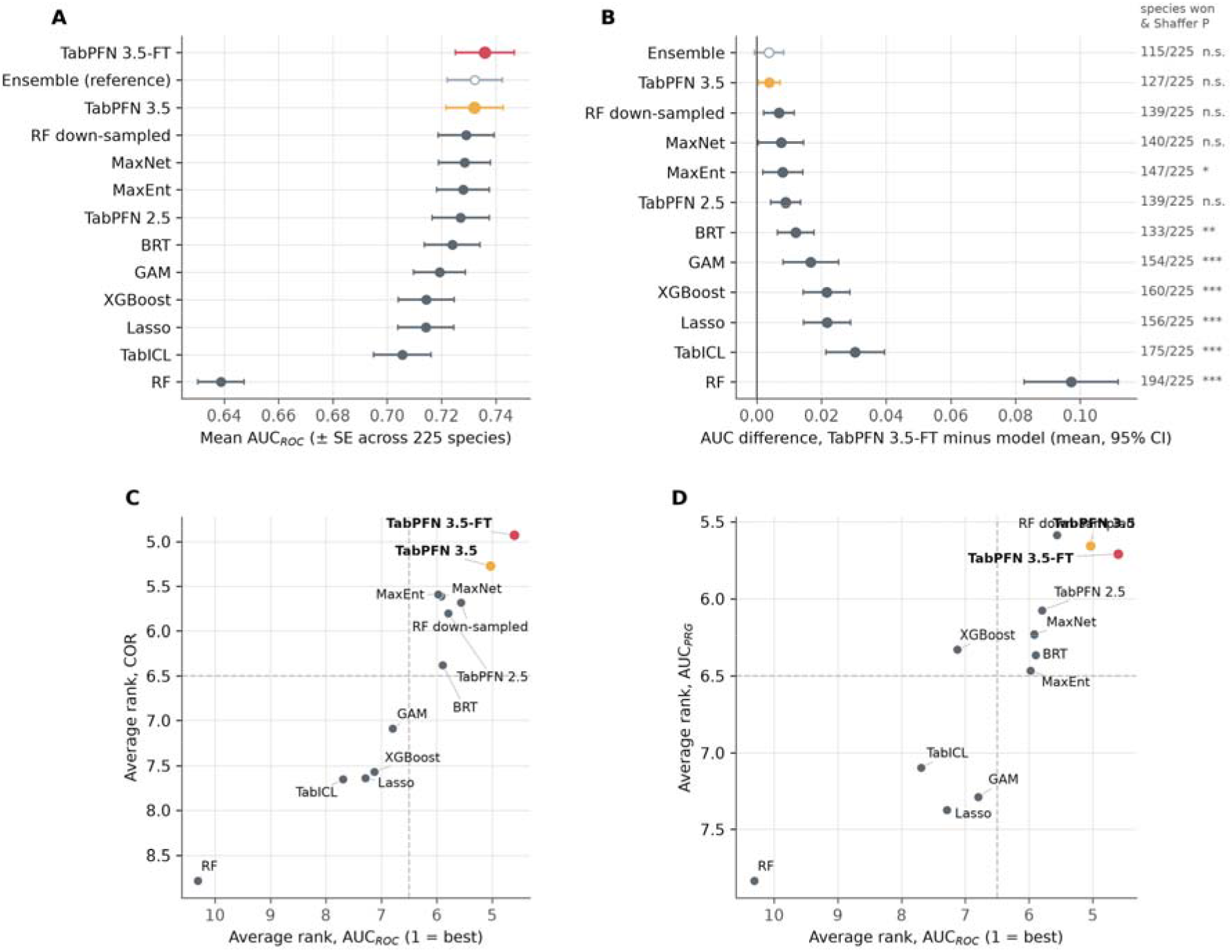
Overall performance on independent presence–absence data (225 species). (A) Mean AUC (± SE across species) of the 12 single models and the five-model ensemble (open symbol; reference). TabPFN 3.5-FT is fine-tuned with leave-one-region-out cross-validation, so its weights never saw the evaluated region. (B) Paired AUC difference between TabPFN 3.5-FT and each model (mean and 95% CI across species; region-level intervals are in Table S5). The numbers give the species for which TabPFN 3.5-FT had the higher AUC and the significance from Shaffer-corrected aligned-rank post hoc tests (* *P*<0.05; ** *P*<0.01; *** *P*<0.001; n.s., not significant). (C, D) Average rank across species of the 12 single models for AUC against COR (C) and against AUC_PRG_ (D); 1 is best, and dashed lines mark the middle rank.

Differences among species dwarfed differences among methods (Fig. 2). In a two-way analysis of the 12 single models, species accounted for 89% of the variation in AUC and models for 2.6%. For the same model, mean AUC ranged from 0.54 in Ontario to 0.82 in Switzerland. Fig. S11 shows predicted distribution maps for one example species.

**Figure 2.**
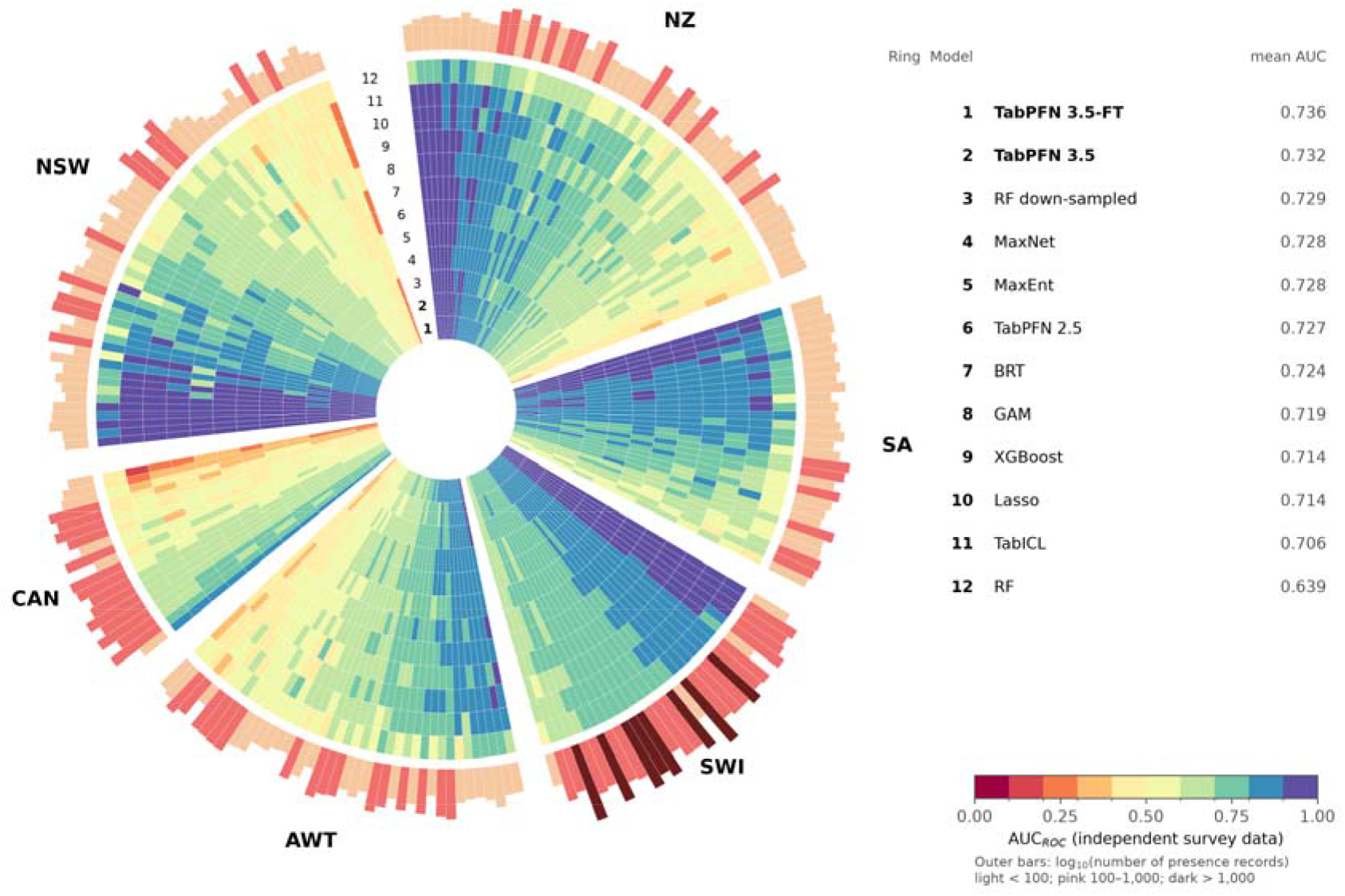
Per-species AUC of 12 single models on independent presence–absence data (225 species). Each spoke is a species and each ring a model; species are grouped by region and sorted by mean AUC within each region. Ring numbers (shown in the gap at the top) follow the key on the right, which lists models by overall mean AUC. The outer bars show the number of presence records per species on a log scale.

### Fine-tuned TabPFN is the best single model on the benchmark

We fine-tuned TabPFN 3.5 on presence–background tasks built from the species of five regions and evaluated it on the sixth, rotating through all regions (TabPFN 3.5-FT); the evaluated region contributed no data to fine-tuning, and all settings were fixed in advance. TabPFN 3.5-FT had the highest mean AUC (0.736), point-biserial correlation (COR, 0.220) and true skill statistic (TSS, 0.446) of all single models, and the best average ranks for these metrics (Fig. 1A, C). With aligned-rank tests and Shaffer’s correction among the 13 approaches, it significantly outperformed MaxEnt (+0.008 in AUC; higher for 147 of 225 species; *P* = 0.021), BRT (+0.012; *P* = 0.005), GAM, Lasso, XGBoost, TabICL and default random forests (*P* < 0.001; Fig. 1B), but not MaxNet (*P* = 0.09), RF down-sampled (*P* = 0.33) or TabPFN 2.5 (*P* = 0.052), and it did not differ from the ensemble (*P* = 1). The gain was not due to how categorical covariates were encoded (mean AUC 0.726 with both encodings, five bags).

These tests treat species as independent, although they are nested in six regions and share survey sites within biological groups. A bootstrap of species within regions gave a 95% interval of 0.002 to 0.014 for the difference from MaxEnt, which was positive in five of six regions, but a mixed model with region as a random effect estimated 0.004 with an interval that included zero (−0.012 to 0.020; Table S5). The significant contrasts also required the 100-bag setting: with five bags, the difference from MaxEnt was 0.005 (−0.002 to 0.011). For AUC_PRG_, TabPFN 3.5-FT had the highest median but not the highest mean, because of one Ontario species with a strongly negative value (Fig. 1D; Table S4).

TabPFN 3.5-FT was the best single model for 17.8% of species, more often than any other (RF down-sampled 10.7%, MaxEnt 5.1%; Fig. S10A), and its advantage over MaxEnt did not change with the number of presence records (*P* = 0.87; Fig. S10B). It exceeded all 21 published methods in four of six regions (Fig. S12) and fell in the lower part of the published range only in Ontario (0.543; range 0.535–0.595), where every method performed worst.

### Fine-tuned TabPFN gives better-calibrated predictions

Presence–background models estimate relative likelihood, so the absolute values of their predictions cannot be compared with survey data. Their calibration slope, however, can be estimated (7). This is the slope of a logistic regression of observed presence at the survey sites on the logit of the prediction. A slope of 1 means that differences in predicted log-odds between sites are right, whereas a slope below 1 means that predictions are too extreme. The median slope was 0.78 for TabPFN 3.5-FT (interquartile range 0.45–1.19), similar to BRT (0.76) and RF down-sampled (0.75), and much closer to 1 than for MaxEnt and MaxNet (both 0.33; Fig. S9). Fine-tuning moved TabPFN 3.5 closer to 1 (from 0.69). Averaging over many bags shrinks predictions and therefore also raises the slope, but this explains only part of the difference: with five bags the slope was 0.72 for the fine-tuned model and 0.65 without fine-tuning, still about twice that of MaxEnt.

### Virtual species: it depends on the relationship and on sampling bias

We simulated 120 virtual species on the real landscapes of the six regions, 60 with additive responses to two or three covariates and 60 with complex, non-additive relationships, each fitted with nested samples of 10 to 300 presence records (Fig. 3; Table S7). With additive relationships, the advantage of the foundation model disappeared. TabPFN 3.5-FT recovered the true probability of occurrence less well than MaxEnt (mean difference in Spearman correlation −0.046; better for 80 of 240 datasets; *P* < 10^−8^), Lasso, GAM and the ensemble, as well as BRT (*P* = 0.29) and better than RF down-sampled, and the regression-based methods recovered response curves better, especially with many records (0.88–0.92 with 300 records, against 0.79). Part of this gap arose from extrapolation: inside the environmental envelope of the presence records, the deficit relative to MaxEnt halved (−0.020) and that relative to GAM disappeared. With complex relationships the gap narrowed, and without sampling bias the untuned foundation model recovered the true probability at least as well as every single model (Spearman correlation 0.71, against 0.70 for MaxNet and 0.69 for MaxEnt; *P* = 0.07 and 0.05), second only to the ensemble (0.72), and it was the best single model for multiplicative interactions. Fine-tuning removed most of this advantage (−0.05 with 100 and 300 records; *P* < 10^−6^). In both designs, the foundation models were clearly worse when records were collected with environmental sampling bias (by 0.05–0.07 relative to MaxEnt; *P* < 10^−4^). This weakness was explained neither by correlation between the bias covariate and the covariates that determine occurrence (*P* = 0.24) nor by the small number of background points in each balanced bag, because five times more background points made recovery worse (−0.038 with 10 records; *P* = 0.002). Against the simulated presences and absences, the foundation model again gave the best-calibrated predictions (median slope 0.96 for TabPFN 3.5-FT, against 0.43 for MaxEnt). Its partial responses usually had the right direction but were attenuated (Fig. S13).

**Figure 3.**
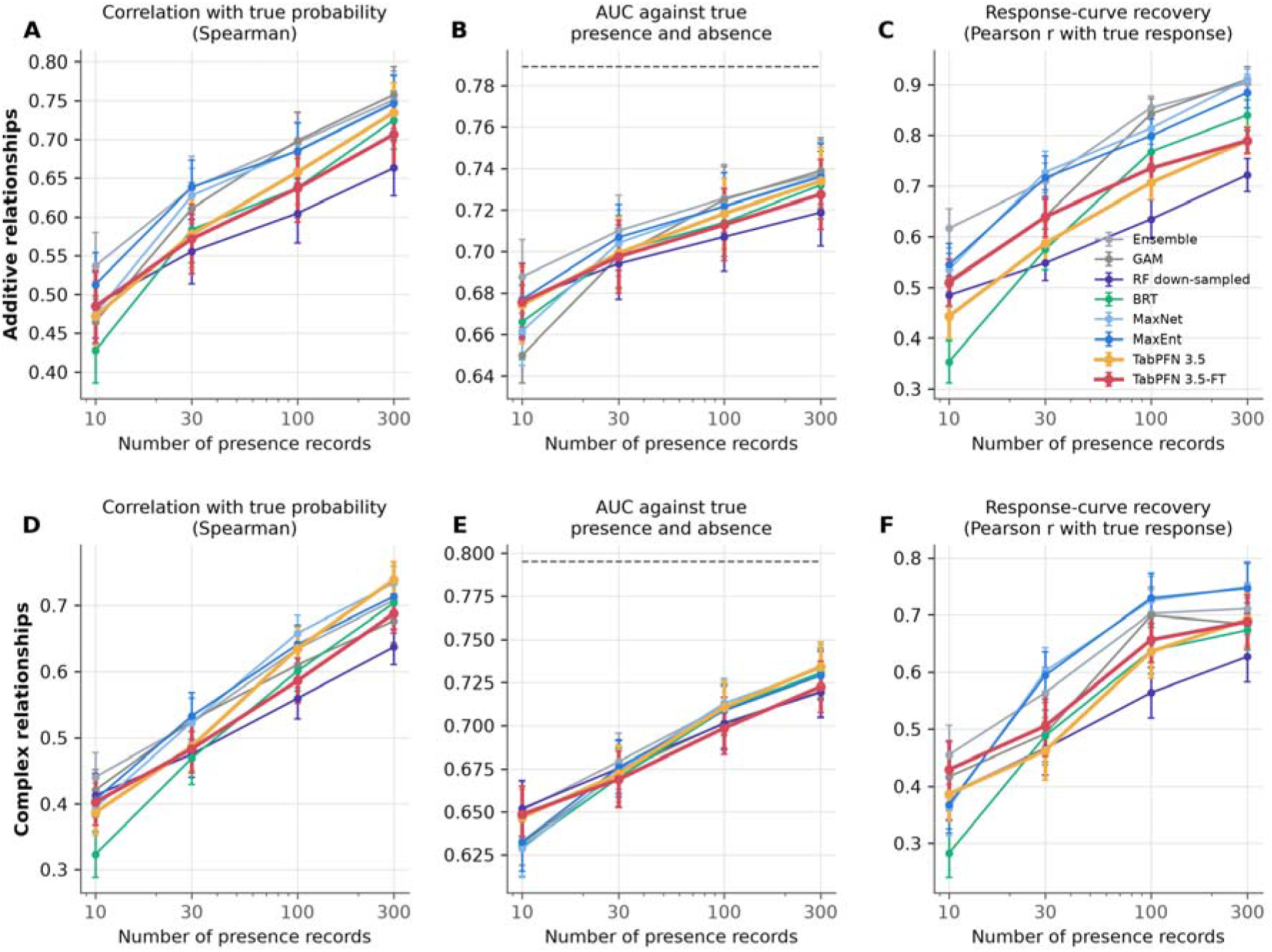
Virtual species with known distributions. Top (A–C): 60 species with additive responses to two or three covariates. Bottom (D–F): 60 species with non-additive relationships. Each species was fitted with 10, 30, 100 and 300 presence records; half had environmental sampling bias. (A, D) Spearman correlation between predicted and true probability of occurrence at 5,000 random landscape points. (B, E) AUC against simulated presences and absences; the dashed line is the AUC of the true probability. (C, F) Pearson correlation between each model’s partial-dependence curve and the true partial response. Points are means ± SE across species; TabPFN used 20 balanced bags.

### What the model needs to learn: presence–background structure, not absences

All fine-tuning strategies were compared with the same leave-one-region-out design and five-bag predictions (Fig. 4; Table S3). The pre-specified strategy, fine-tuning on presence– background tasks, raised mean AUC from 0.726 to 0.733. Supervision with the presence– absence surveys of the training regions did not help (0.727 after 10 epochs; 0.726 with the number of epochs chosen by nested cross-validation). Virtual species, which need no field data, gave 0.730 alone and 0.734–0.735 when mixed with real tasks. Gains were concentrated in the Australian Wet Tropics and New Zealand; most strategies slightly lowered AUC in Switzerland, and none improved Ontario. The generic prior thus gains most from learning the structure of presence–background tasks rather than what absences look like.

**Figure 4.**
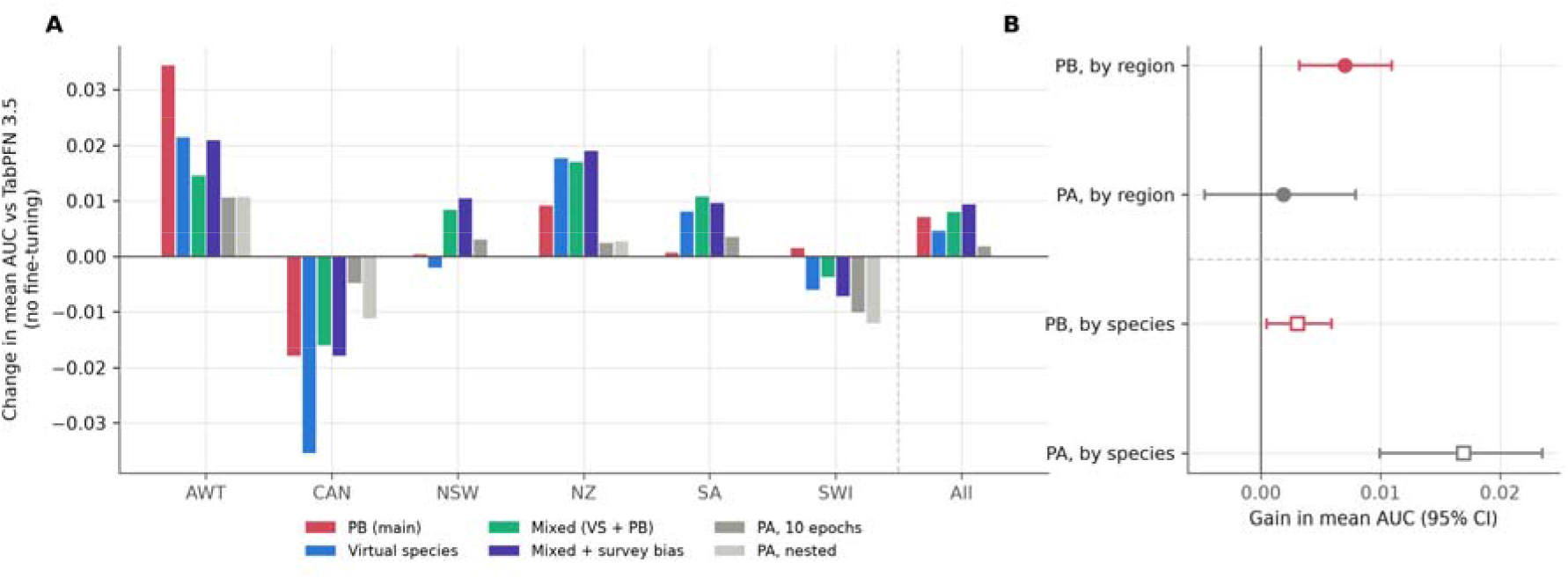
Fine-tuning strategies. (A) Change in mean AUC relative to TabPFN 3.5 without fine-tuning, in each held-out region and overall. Each region was predicted by a model fine-tuned only on the other five regions. (B) Gain in mean AUC (95% CI across species) when fine-tuning held out whole regions (circles) or used other species from all regions, including the evaluated ones (squares). All arms use the same five-bag predictions and seeds. PB, presence–background tasks (main strategy); PA, presence–absence tasks; VS, virtual species.

When fine-tuning instead used other species from all regions, including the evaluated ones (a species-level split, as in the previous study (34)), presence–absence supervision raised mean AUC by 0.017 (to 0.743), against 0.002 when the evaluated region was held out (difference *P* < 10^−11^; Fig. 4B); with such data it even improved Ontario (+0.034). Presence–background tasks gave similar gains under both designs (+0.003 and +0.007; *P* = 0.33). The evaluated species’ own labels were never used, so the larger gain reflects surveys of other species at the same sites. It may come from better predictions across the region or partly from properties of the surveyed sites themselves; the benchmark cannot separate the two.

### Performance holds under spatially separated evaluation

Under the spatial-block design of Valavi et al. (38), in which models were trained on presence records with target-group background and tested at survey sites in held-out blocks (178 species), spatial separation lowered mean AUC by similar amounts for all models (0.016 for TabPFN 3.5-FT, 0.014–0.017 for the main reference methods; Fig. S14). The ensemble ranked first (0.722), TabPFN 3.5-FT second (0.721), and MaxNet and MaxEnt followed (0.718); after Shaffer’s correction, TabPFN 3.5-FT differed significantly only from default random forests, and its advantage over MaxEnt was small (+0.003; *P* = 0.46). Fine-tuning still improved TabPFN 3.5 (+0.008; *P* = 0.005). With target-group background, Ontario was no longer an outlier (0.715 against 0.543 in the main benchmark) and fine-tuning no longer lowered AUC there, suggesting that the Ontario deficit reflects sampling bias that target-group background partly corrects. These analyses used five bags, so the comparison is conservative for TabPFN.

## Discussion

A tabular foundation model, TabPFN 3.5, used without any tuning, performed as well as the best of 21 established SDM methods on the standard benchmark for presence-only prediction, and after fine-tuning on presence–background tasks from other regions it became the best single model there. The gains over MaxEnt were modest (+0.008 AUC) but consistent across species and five of six regions, and came with better-calibrated predictions. With only six regions, however, the benchmark cannot show that the advantage holds in new regions.

The virtual species show where the foundation model helps and where it does not. With simple additive relationships, the structure assumed by GAM, Lasso and MaxEnt, these methods recovered the truth better, as simulations favour methods whose structure matches the simulated one (36, 39); with more complex relationships and unbiased records, the untuned foundation model was at least as good as any single model. Its clearest weakness was sensitivity to environmental sampling bias, whose mechanism remains unclear, and fine-tuning traded part of its flexibility for better discrimination at survey sites. Its calibration advantage held in every setting. Consistent with the role of bias, target-group background removed both the poor performance in Ontario and the loss from fine-tuning there.

The fine-tuning experiments show what the generic prior lacks. Supervision with true absences from other regions did not help, but presence–background tasks and simulated species did, so the default prior seems poorly matched to the structure of presence– background data rather than to the shape of species’ responses. Our species-split experiment helps to explain why the only previous study reported a much larger gain from fine-tuning, from 0.728 to 0.762 in mean AUC (34): presence–absence supervision raised mean AUC by 0.017 when it came from other species of the evaluated regions, but by only 0.002 when it did not. The two designs answer different questions. Leave-one-region-out estimates the gain in a region without surveys, the usual situation for presence-only data. The species split estimates the gain where other species have already been surveyed at the same sites; fine-tuning on such surveys may then be worthwhile, but its gain is measured at the very sites used for fine-tuning. Evaluations of fine-tuned SDMs should state which situation they represent.

If they generalise, these results matter beyond ecology. MaxEnt and related methods are default tools for presence-only prediction in conservation, invasion biology, epidemiology, marine science, paleobiology and archaeology (10–12, 15–17), often with default settings (24). A model that matches MaxEnt at survey sites with fixed settings and better calibration could be a useful default for ranking sites, for example to target surveys. Where the aim is to describe a niche, or records are environmentally biased, its response curves should be compared with those of a regression-based method, and bias should be corrected, for example with target-group background.

Several limitations apply. The benchmark has 225 species in six regions, with records collected decades ago, and the results must be tested on other taxa, regions and kinds of presence-only data. Evaluation at survey sites measures discrimination, not transfer to new environments; under spatial-block cross-validation the fine-tuned model lost no more accuracy than established methods, but its advantage over MaxEnt was not significant. Our virtual species represent one simple class of niches. TabPFN weights are released under a non-commercial licence, and the 100-bag setting takes about 2 min per species on one GPU, against about 20 s for MaxEnt (Table S6). Finally, if a few hundred real presence– background tasks improve a generic tabular prior, a model pre-trained on many thousands of species with realistic observation processes could become a foundation model for species distributions; we are building such a model for North American birds.

## Materials and methods

### Benchmark data

We used the NCEAS benchmark data (20, 21) in exactly the form used by Valavi et al. (22). The data cover 225 species in six regions: the Australian Wet Tropics (AWT; 40 species of birds and plants), Ontario, Canada (CAN; 20 birds), New South Wales (NSW; 53 species in eight biological groups), New Zealand (NZ; 52 plants), South America (SA; 30 plants) and Switzerland (SWI; 30 trees). For each species, presence-only records from museums, herbaria and atlases serve as training data (5–5,822 records; median 57). Independent presence–absence surveys serve as test data, with 102 to 19,120 sites per species. All species in a biological group share the same survey sites.

Following Valavi et al. (22), we used for each region the set of covariates with pairwise correlations of at most 0.8 (7–12 covariates; Table S1). Categorical covariates (vegetation type in CAN and NSW, age class and soil toxicity in NZ, and calcareous substrate in SWI) were treated as factors. Geographic coordinates were not used as predictors. For each region we drew 50,000 random background points from the covariate rasters (seed 42), and all models used the same points. Continuous covariates were standardised with the mean and standard deviation of each species’ training data (presences plus background points). The same transformation was applied to the test sites.

### Reference models

We re-fitted nine methods with the settings and code of Valavi et al. (22) and added XGBoost (Table 1); the remaining 12 of the 21 published methods were taken from their published results. The reference models span the main families of SDM methods: regression models with smooth or penalised terms (GAM (40) and Lasso (41)), maximum-entropy models (MaxEnt (19, 42) and MaxNet (43)), boosted trees (BRT (44) and XGBoost (45)), random forests with and without down-sampling (46), and the five-model ensemble that performed best in the original benchmark. Where appropriate, background points were down-weighted so that presences and background had equal total weight. Tuning parameters were chosen by internal cross-validation where the original protocol did so (Lasso penalty, number of boosted trees). Pre-specified fallbacks were applied when a model failed to converge. Each model, its settings and the fallbacks are described in detail in the Supplementary Methods (Section S2).

**Table 1.**
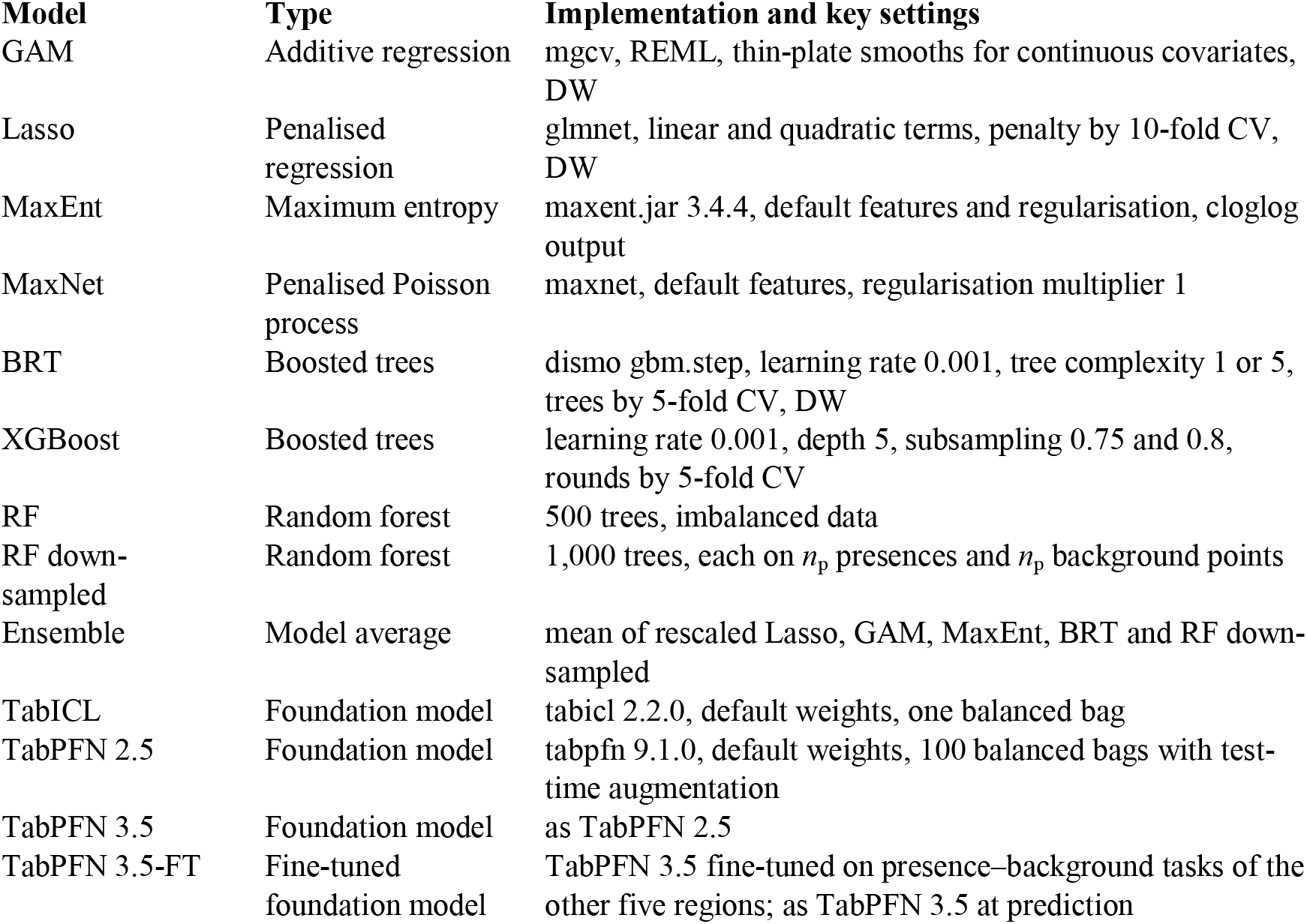
Models compared in this study. All were fitted to the same presence and background data for each species. DW, down-weighting of background points to equal total class weight.

### Foundation models and balanced bagging

TabPFN 3.5 (33) and TabPFN 2.5 (32) were run with the tabpfn package 9.1.0, and TabICL (37) with tabicl 2.2.0. All three used their default weights and hyperparameters. These models receive the training table as context and output class probabilities for the query points in a single forward pass. Their pre-training tasks rarely contain the extreme imbalance of presence–background data. We therefore presented them with balanced bags. Each bag contained the *n*_p_ presences, resampled with replacement, and *n*_p_ points drawn at random from the 50,000 background points. The model predicted the test sites from each bag, and predictions were averaged across bags on the logit scale. For TabPFN, the main results used 100 bags with test-time augmentation (“Stable” setting). In each bag, the test sites were predicted both as observed and with Gaussian noise (SD 0.01 on the standardised scale) added to their continuous covariates, and both predictions entered the average. Comparisons of fine-tuning strategies used 5 bags without augmentation (“Fast” setting). TabICL was run with a single balanced bag. Before the main analysis, we compared eight alternative bagging configurations on half of the species, varying the number of background points per bag, the bootstrap and the number of estimators. None improved on the simple balanced rule (Table S2).

### Fine-tuning

We fine-tuned TabPFN 3.5 with leave-one-region-out cross-validation: for each region, a separate model was fine-tuned only on the species of the other five regions and then used to predict that region. In each epoch, every training species contributed a new class-balanced sample of up to 2,000 presences and as many random background points, which TabPFN’s fine-tuning loop split into context (70%) and query (30%) sets; the loss was the cross-entropy of the query labels, so the network learns the task it faces at prediction time. We used AdamW (learning rate 10^−5^, weight decay 0.01) for 10 epochs. Categorical covariates were coded numerically in fine-tuning and at prediction; with default weights this coding gave the same mean AUC as categorical features (0.726). All settings were fixed before any evaluation, and this presence–background strategy was designated as the main analysis before any other was tried.

We compared it with presence–absence supervision (query sets formed by the survey sites of training-region species, for 10 epochs or a nested-selected number of epochs), fine-tuning on 1,000 simulated species per epoch, mixtures of simulated and real tasks (with or without a survey-effort bias surface (6)) and an ensemble of fine-tuned models (Supplementary Information). To measure the effect of using species from the evaluated regions, we repeated the presence–background and presence–absence strategies with a species-level split, as in the previous study (34): species of each region were assigned at random to five folds, and each fold was predicted by a model fine-tuned on the other four folds from all regions, with all other settings and seeds unchanged.

### Virtual species

To ask whether models recover the true distribution, we simulated 60 virtual species (35, 36, 47), ten on the landscape of each region, represented by its 50,000 background points. The design was fixed before any model was run. Each species responded additively, on the logit scale, to two or three continuous covariates; each response was unimodal (Gaussian) with probability 0.7 and monotone otherwise. The intercept was set to give a prevalence between 0.03 and 0.30, and true occurrence was drawn at every point. Other covariates were irrelevant but were given to all models. Each landscape was split into 5,000 test points, 20,000 background points and the remainder. Presences were drawn from occupied points outside the test set, uniformly for half of the species and with environmental sampling bias on a covariate outside the niche for the other half, as nested samples of 10, 30, 100 and 300 records (240 datasets). All reference models and TabPFN 3.5, with default and leave-one-region-out fine-tuned weights (20 balanced bags), were fitted with the benchmark settings. We measured the Spearman correlation between predicted and true probability of occurrence, AUC against the simulated presences and absences, and the recovery of response curves: the Pearson correlation between each model’s partial-dependence curve (mean logit over 100 test points with one niche covariate set to each of 12 quantiles) and the true additive response, averaged over niche covariates. A second set of 60 species, simulated after the first had been analysed, had non-additive relationships (correlated tolerances, joint thresholds or multiplicative interactions) with otherwise identical settings. We also recomputed the Spearman correlation inside the environmental envelope of the presences, and tested whether, among biased species, the deficit of the foundation model increased with the correlation between the bias covariate and the niche covariates. Details are in the Supplementary Information.

### Evaluation

All models were evaluated on the independent presence–absence surveys. As in Valavi et al. (22), we computed five metrics. The area under the receiver operating characteristic curve (AUC) measures how well a model ranks presences above absences. The area under the precision–recall–gain curve (AUC_PRG_) (48) emphasises performance on presences; we computed it with a Python port of the R prg package that reproduced its output exactly. The point-biserial correlation (COR) between predictions and observations also reflects how far apart the predictions for presences and absences are. The maximum true skill statistic (TSS) is the largest value of sensitivity plus specificity minus one over all thresholds. Average precision summarises the precision–recall curve. A model returning a constant prediction was scored AUC 0.5, COR 0 and TSS 0.

### Statistical analysis

As in Valavi et al. (22), we compared models with the Friedman aligned-ranks test (49, 50), with species as blocks, followed by pairwise post hoc tests with Shaffer’s correction (51), using a Python port of scmamp (52) that reproduced its results exactly. The comparison family comprised the 13 approaches in Fig. 1A (the 12 single models, with 100 bags for the foundation models, and the ensemble). Because species are nested in regions and share survey sites within biological groups, we also computed a bootstrap of species within regions (5,000 resamples) and a linear mixed model of paired differences with region as a random effect. Dependence of the advantage on sample size was tested by regressing paired differences on log_10_ of the number of presences, with region as a fixed effect. Calibration was measured by the calibration slope, the slope of a logistic regression of observed presence on the logit of the prediction, which does not depend on the unknown prevalence (7, 53).

### Spatially separated evaluation

Following Valavi et al. (38), each species was trained on its presence records plus the target-group background of its biological group, using their published samples. Training records and survey sites were assigned to their published spatial blocks, and blocks were allocated at random to 5 folds. We kept the most even of 100 random allocations, judged by the smallest number of records of any class in any fold and then by the standard deviation of these numbers. Fold *f* was trained on the records outside *f* and tested on the survey sites inside *f*. Random 5-fold cross-validation used the same numbers of training and test records. A species was excluded if any fold lacked a class, if any training fold had fewer than 5 presences, or if it had fewer than 10 test presences in total. This left 178 species and 1,780 fold tasks. As in Valavi et al. (38), the vegetation-type covariate was not used in NSW. AUC was computed for each fold and averaged per species. TabPFN 3.5-FT used the leave-one-region-out checkpoints, so no checkpoint had seen data from the region it predicted.

### Computation

R models (R 4.5.2) were fitted on CPU, one core per species with 12 species in parallel. Foundation models and fine-tuning ran on a single NVIDIA GB10 GPU (DGX Spark; PyTorch 2.14.1). Run times are given in Table S6.

## Acknowledgments

This work was supported by JST CREST, Japan (Grant Number: JPMJCR24J2) and the Environment Research and Technology Development Fund of the Environmental Restoration and Conservation Agency (ERCA; Grant Number: JPMEERF26S21300), funded by the Ministry of the Environment, Japan. The funders had no role in study design, data collection and analysis, decision to publish, or preparation of the manuscript.

## Supplementary Material

Supplementary material is available at *PNAS* online.

## Author Contributions

H.D. conceived the study, designed the analytical framework, wrote the analysis code, performed the analyses and wrote the first draft of the manuscript. T.O. and N.T. contributed to the study design and to the ecological and methodological interpretation of the results. All authors interpreted the results, revised the manuscript critically and approved the final version.

## Data Availability

The NCEAS benchmark data are openly available (21). The spatial blocks and target-group background samples are from Valavi et al. (38). All code, per-species predictions and results are deposited on GitHub (https://github.com/hideyuki-doi/tabfm-distribution). TabPFN 3.5 weights are under the Prior Labs non-commercial licence.

## Artificial Intelligence

Claude (Anthropic) was used to help write and debug the analysis code, run the analyses, edit the manuscript text, and check the references. The authors reviewed and verified all AI-assisted outputs and take full responsibility for the content of the manuscript.

