## Supplemental materials for "A tabular foundation model rivals established species distribution models from presence-only records"

**This PDF file includes:**

Supplementary text

Figures S1 to S14

Tables S1 to S7

SI References

**Supplementary Information Text**

**S1. Data and preprocessing**

We used the NCEAS data (1, 2) exactly as distributed for the benchmark of Valavi et al. (3). The data provide presence-only training records and independent presence–absence test sites for 225 species: 40 in the Australian Wet Tropics (AWT), 20 in Ontario, Canada (CAN), 53 in New South Wales (NSW), 52 in New Zealand (NZ), 30 in South America (SA) and 30 in Switzerland (SWI). The covariates are the weakly correlated sets (|*r*| ≤ 0.8) defined by Valavi et al. in their code (nceas_model_vars.R) and listed in Table S1. Categorical covariates were treated as factors in the R models and were one-hot encoded for XGBoost. TabPFN received them as categorical features, except in the fine-tuning experiments, where they were coded numerically (Section S4).

For each region we drew 50,000 background points at random from the covariate rasters (seed 42), and every model used the same points. Geographic coordinates were not used as predictors. Continuous covariates were z-scored with the mean and standard deviation of each species’ training data (presences plus background points), and the same transformation was applied to its test sites.

**Table S1.** Covariates used in each region. Categorical covariates are marked with an asterisk.

| **Region** | **Covariates** |
| --- | --- |
| AWT | bc04, bc05, bc06, bc12, bc15, slope, topo, tri |
| CAN | alt, asp2, ontprec, ontslp, onttemp, ontveg*, watdist |
| NSW | cti, disturb, mi, rainann, raindq, rugged, soildepth, soilfert, solrad, tempann, topo, vegsys* |
| NZ | age*, deficit, hillshade, mas, mat, r2pet, slope, sseas, toxicats*, tseas, vpd |
| SA | sabio2, sabio4, sabio5, sabio6, sabio12, sabio15, sabio17, sabio18 |
| SWI | bcc, calc*, ccc, ddeg, nutri, pday, precyy, sfroyy, slope, sradyy, swb, topo |

**S2. Reference models**

We re-fitted nine methods with the settings and code of Valavi et al. (3) and added XGBoost (Table 1 of the main text). The remaining 12 of the 21 published methods were taken from their published results. Several models used down-weighting, in which presences receive weight 1 and background points weight *n*_p_/*n*_b_, so that the two classes have equal total weight.

Generalised additive models (GAM) were fitted with mgcv (4) using restricted maximum likelihood (REML), a binomial logit link and down-weighting. Continuous covariates with at least 15 distinct values entered as thin-plate regression splines with default basis dimension, other covariates as linear terms, and factors as parametric terms. The Lasso was a logistic regression on orthogonal linear and quadratic terms of each covariate, fitted with glmnet (5) with down-weighting. The penalty was chosen by 10-fold cross-validation (λ minimising deviance).

Boosted regression trees (BRT) were fitted with the gbm.step procedure of dismo and gbm (6), using Bernoulli loss and down-weighting. Tree complexity was 1 for species with fewer than 50 presences and 5 otherwise, the learning rate was 0.001 and the bag fraction 0.75. The number of trees (at most 10,000, in steps of 50) was chosen by 5-fold cross-validation. XGBoost (7) used a learning rate of 0.001, maximum depth 5, row subsampling 0.75 and column subsampling 0.8. Its number of rounds (at most 15,000) was chosen by stratified 5-fold cross-validation on AUC.

Default random forests (RF) were classification forests of 500 trees grown with randomForest (8) on the full, imbalanced data. Down-sampled random forests (RF down-sampled) grew 1,000 trees, each on a bootstrap sample of *n*_p_ presences and *n*_p_ background points. This balances the classes within each tree without discarding background information across the forest.

MaxEnt estimates the distribution of maximum entropy over the background, subject to the constraint that the expected values of the covariate features match their means at the presences (9, 10). We used maxent.jar 3.4.4 with default settings: automatic feature classes (linear, quadratic, product and hinge features, enabled according to the number of presences) and the default regularisation multiplier of 1. Categorical covariates were declared as categorical layers, and we used the cloglog output. MaxNet (11) fits the same model as an L1-penalised inhomogeneous Poisson process through glmnet, with default features and a regularisation multiplier of 1.

The ensemble averaged the predictions of Lasso, GAM, MaxEnt, BRT and RF down-sampled, each rescaled to the range 0–1. This combination performed best in the original benchmark (3). When a model failed to converge, we applied pre-specified fallbacks, all fixed before evaluation. A GAM with factor terms that had not converged after 30 min was refitted without them, as Valavi et al. specified for NZ and CAN. Lasso runs that did not converge were repeated with more iterations and a shorter regularisation path. BRT was retried with smaller learning rates. MaxNet was retried with a relaxed convergence threshold.

The software versions and the exact fallback rules are as follows. We used mgcv 1.9-1, glmnet, the gbm.step source of dismo 1.3-14 with gbm 2.3.1, randomForest 4.7-1.2, maxnet 0.1.4, maxent.jar 3.4.4 and xgboost 3.4.1. MaxEnt was run with the “nothreshold” option, so that threshold features were not used, and with categorical layers declared as categorical. XGBoost also used a minimum child weight of 1 and γ = 0, without case weights. In the ensemble, a member that returned a constant prediction contributed 0 after rescaling.

All fallback rules were fixed before evaluation. Each model had a time limit of 3 h. A GAM with factor terms that had not converged after 30 min was refitted without them; this is how Valavi et al. (3) specified their GAMs for NZ and CAN. If cv.glmnet failed to converge at small values of λ, the Lasso was refitted with maxit =10^6^ and lambda.min.ratio =0.01. If gbm.step found that cross-validated deviance did not improve from the first step, BRT was refitted with learning rates of 5×10^−4^ and then 10^−4^, and finally with 10^−4^ and a step of 10 trees. If all attempts failed, the optimal BRT had no trees, and the intercept-only prediction was used. If glmnet failed to converge inside MaxNet, the same model was refitted with a convergence threshold of 10^−5^; if that also failed, we used the regularisation path up to the last λ that had converged. A model that returned a constant prediction was scored AUC =0.5, COR =0 and TSS =0.

Fig. S1 compares our regional mean AUCs with the published values. The mean absolute difference was 0.001–0.006 for eight methods and 0.013 for default random forests.

**S3. Foundation models and balanced bagging**

TabPFN 3.5 and 2.5 (12, 13) were run with tabpfn 9.1.0, and TabICL (14) with tabicl 2.2.0. All used default weights and hyperparameters (PyTorch 2.14.1, CUDA 13). In each bag, the *n*_p_ presences were resampled with replacement and *n*_p_ points were drawn at random from the 50,000 background points. The model was given this balanced table as context and predicted the test sites. Predictions were averaged across bags on the logit scale. The “Stable” setting used 100 bags with test-time augmentation: in each bag the test sites were predicted both as observed and with Gaussian noise (SD 0.01 on the z-scale) added to their continuous covariates, and both predictions entered the average. The “Fast” setting used 5 bags without augmentation. TabICL was run with a single balanced bag.

Before the main analysis, we compared eight bagging configurations on a random half of the species in each region (the development set). Results are for the 73 species completed before the search was stopped (Table S2). No configuration improved on balanced bags with bootstrapped presences. Adding background points or estimators reduced AUC, so the search was not used further.

**Table S2.** Bagging configurations for TabPFN 3.5 (development species, *n* = 73; 10 bags unless stated).

| **Configuration** | **Background per bag** | **Mean AUC** |
| --- | --- | --- |
| Presences bootstrapped (default) | *n*_p_ | 0.710 |
| All presences, no bootstrap | *n*_p_ | 0.707 |
| All presences, 16 estimators | *n*_p_ | 0.705 |
| All presences | 3*n*_p_ | 0.705 |
| All presences, 16 estimators | 3*n*_p_ | 0.706 |
| All presences | 10*n*_p_ | 0.701 |
| All presences | max(*n*_p_, 2000) | 0.694 |
| All presences (3 bags) | max(*n*_p_, 10000) | 0.687 |

**S4. Fine-tuning**

**Implementation.** We used FinetunedTabPFNClassifier from tabpfn 9.1.0, with AdamW (learning rate 10^−5^, weight decay 0.01), two estimators per step, no validation split and no early stopping. By default, the fine-tuning loop draws context and query splits from a single training table. We replaced its data-chunking function so that each epoch supplied one task per training species, and a custom split function returned each task’s own context and query sets. Categorical covariates were passed as numeric codes, because the loop uses one categorical setting for all tasks. The comparison arm without fine-tuning used the same coding, so every comparison in Fig. 4 and Table S3 is like-for-like. All settings, including the number of epochs, were fixed before any evaluation on held-out regions. Each held-out region was predicted only by a model fine-tuned on the other five regions.

**Presence–background tasks (PB; main strategy).** In each epoch, every species in the training regions contributed one balanced sample: all of its presences (at most 2,000) and the same number of random background points. The loop split each sample into context (70%) and query (30%), and the loss was the cross-entropy of the query labels. Training ran for 10 epochs. This strategy needs nothing beyond the presence–background data that are available for any species.

**Presence–absence supervision (PA).** The context was a species’ balanced presence–background sample, as above. The query was up to 2,000 of that species’ independent survey sites, with at most half of them presences, labelled with the observed presence or absence. Only the surveys of the training regions were used. This strategy therefore uses information that the published methods did not have. We ran it for 10 epochs and, separately, with the number of epochs chosen by nested cross-validation. For each outer region *R* and each pair of regions {*R*, *Q*}, we fine-tuned on the remaining four regions and scored epochs 0, 1, 2, 3, 5, 7 and 10 on *Q* (2 bags, at most 3,000 test sites per species). For *R* we chose the epoch with the best mean AUC over the five regions *Q* ≠ *R*. The chosen epochs were 5 for AWT, 3 for CAN, 0 for NSW, 3 for NZ, 0 for SA and 3 for SWI.

**Virtual species (VS).** The landscapes were the 50,000 background points of the training regions, with their real covariate values and coordinates. In each of 5 epochs, 1,000 new species were simulated, each on a randomly chosen training region. Each species responded to 1–4 continuous covariates. A response was unimodal (Gaussian, with its optimum in [−1.5, 1.5] and a width log-uniform in [0.3, 2]) with probability 0.7, and monotone (slope 0.5–3) otherwise. With probability 0.3, random effects of one categorical covariate were added. The linear predictor was standardised and multiplied by a random steepness of 1–4, and the intercept was set to give a prevalence log-uniform in 1–40%. True occurrence was then drawn from a Bernoulli distribution at every landscape point. Sampling intensity was uniform with probability 0.25, environmental ((±β *x*) for one covariate, with β in [0.5, 2.5]) with probability 0.35, and spatial (Gaussian kernels around 1–5 random hubs, with a bandwidth of 5–30% of the regional extent) with probability 0.40. Between 5 and 1,000 presence records (log-uniform, at most the number of occupied points) were drawn from the occupied points in proportion to sampling intensity. The context was these presences plus the same number of uniform background points, and the query was 1,000 uniform sites labelled with true occurrence.

**Mixed and mixed with target-group bias.** “Mixed” added the real PB tasks of the training regions to the virtual species in every epoch, for 5 epochs. “Mixed + survey bias” also used a real survey-effort surface for each training region. This was a Gaussian kernel density of the pooled presence-only records of all its species (the target-group idea of Phillips et al. (15)), with a bandwidth of 3% of the regional extent. The surface set the sampling intensity of 40% of the virtual species, and this strategy ran for 10 epochs.

**Species-split fine-tuning.** To measure how fine-tuning on species from the evaluated regions changes the gain, the species of each region were assigned at random to five folds (seed 2026), and each fold was predicted by a model fine-tuned for 10 epochs on the other four folds from all six regions. The PB and PA strategies were run in this way with all other settings and evaluation seeds unchanged. Because species of a biological group share survey sites, the PA variant sees the survey sites of the evaluated species, with the labels of other species, during fine-tuning.

**Ensemble of fine-tuned models.** The rule was fixed in advance: average, on the logit scale, five bags from each of the PB, VS and Mixed models (15 bags in total). The control was the PB model with 15 bags, so that a gain from combining models is not confused with a gain from more bags.

**Table S3.** Mean AUC of fine-tuning strategies by held-out region (Fast setting, 5 bags unless stated; categorical covariates as numeric codes). The last two rows use a species-level split, in which the evaluated species’ regions, and their survey sites, were also used for fine-tuning.

| **Strategy** | **AWT** | **CAN** | **NSW** | **NZ** | **SA** | **SWI** | **All** |
| --- | --- | --- | --- | --- | --- | --- | --- |
| No fine-tuning (default) | 0.661 | 0.558 | 0.724 | 0.740 | 0.809 | 0.816 | 0.726 |
| PB, 10 epochs (main) | 0.696 | 0.540 | 0.725 | 0.749 | 0.810 | 0.818 | 0.733 |
| PA, 10 epochs | 0.672 | 0.553 | 0.727 | 0.743 | 0.813 | 0.806 | 0.727 |
| PA, nested epochs | 0.672 | 0.547 | 0.724 | 0.743 | 0.809 | 0.804 | 0.726 |
| VS, 5 epochs | 0.683 | 0.523 | 0.722 | 0.758 | 0.817 | 0.810 | 0.730 |
| Mixed (VS + PB), 5 epochs | 0.676 | 0.542 | 0.733 | 0.757 | 0.820 | 0.812 | 0.734 |
| Mixed + survey bias, 10 epochs | 0.682 | 0.540 | 0.735 | 0.759 | 0.819 | 0.809 | 0.735 |
| PB, 15 bags (control) | 0.699 | 0.538 | 0.725 | 0.753 | 0.806 | 0.820 | 0.734 |
| Ensemble PB + VS + Mixed (15 bags) | 0.688 | 0.537 | 0.729 | 0.758 | 0.818 | 0.817 | 0.735 |
| PB, species split, 10 epochs | 0.672 | 0.542 | 0.726 | 0.749 | 0.811 | 0.815 | 0.729 |
| PA, species split, 10 epochs | 0.695 | 0.592 | 0.736 | 0.754 | 0.821 | 0.821 | 0.743 |

**S5. Evaluation and statistics**

All metrics were computed on the independent presence–absence sites. AUC_PRG_ (16) was computed with a Python port of the R prg package, and COR is the Pearson correlation between predictions and observed presence or absence. Models were compared with the Friedman aligned-ranks test (17, 18), using species as blocks. For pairwise post hoc tests, the statistic was *z* = (*R̄_i_* − *R̄_j_*)/√(*k*(*nk*+1)/6), where *R* are mean aligned ranks, *k* is the number of models and *n* the number of species. The *P* values were corrected with Shaffer’s static procedure (19). Our implementation is a port of scmamp (20), and both ports reproduced the original packages exactly.

The main text compares the 13 approaches in its Fig. 2A, using the Stable setting for the foundation models. Figs. S2–S4 add the Fast variants and the ensembles that include TabPFN. Adding these to the comparison family changes the corrected *P* values: for example, TabPFN 3.5-FT against MaxNet gives *P* = 0.03 among 18 approaches but *P* = 0.09 among 13. Ensembles that include TabPFN combined their members after rescaling, as for the published ensemble.

**All metrics.** Table S4 gives the mean and median of all five metrics for the 13 approaches of the main comparison. AUC_PRG_ is unbounded below, and its mean is strongly affected by a few species with very negative values; the median is more robust.

**Table S4.** Mean / median of each metric across 225 species, independent presence–absence data.

| **Model** | **AUC** | **AUC_PRG_** | **COR** | **TSS** | **Average precision** |
| --- | --- | --- | --- | --- | --- |
| TabPFN 3.5-FT | 0.736 / 0.764 | 0.126 / 0.647 | 0.220 / 0.219 | 0.446 / 0.441 | 0.281 / 0.264 |
| TabPFN 3.5 | 0.732 / 0.755 | 0.142 / 0.643 | 0.213 / 0.204 | 0.438 / 0.442 | 0.278 / 0.257 |
| Ensemble | 0.732 / 0.749 | 0.360 / 0.612 | 0.216 / 0.203 | 0.437 / 0.422 | 0.275 / 0.245 |
| RF down-sampled | 0.729 / 0.738 | 0.393 / 0.623 | 0.211 / 0.207 | 0.431 / 0.411 | 0.276 / 0.238 |
| MaxNet | 0.728 / 0.738 | 0.343 / 0.623 | 0.213 / 0.206 | 0.424 / 0.407 | 0.263 / 0.230 |
| MaxEnt | 0.728 / 0.738 | 0.343 / 0.621 | 0.211 / 0.208 | 0.423 / 0.404 | 0.263 / 0.226 |
| TabPFN 2.5 | 0.727 / 0.743 | 0.264 / 0.622 | 0.208 / 0.200 | 0.435 / 0.430 | 0.275 / 0.249 |
| BRT | 0.724 / 0.740 | 0.316 / 0.613 | 0.207 / 0.189 | 0.429 / 0.418 | 0.265 / 0.235 |
| GAM | 0.719 / 0.717 | 0.176 / 0.528 | 0.196 / 0.184 | 0.418 / 0.397 | 0.245 / 0.221 |
| Lasso | 0.714 / 0.727 | 0.167 / 0.540 | 0.190 / 0.181 | 0.414 / 0.409 | 0.248 / 0.211 |
| XGBoost | 0.714 / 0.722 | 0.367 / 0.600 | 0.179 / 0.151 | 0.408 / 0.392 | 0.265 / 0.231 |
| TabICL | 0.706 / 0.705 | 0.266 / 0.563 | 0.182 / 0.173 | 0.393 / 0.374 | 0.253 / 0.220 |
| RF | 0.639 / 0.612 | 0.297 / 0.474 | 0.132 / 0.090 | 0.265 / 0.215 | 0.223 / 0.188 |

**Uncertainty at the regional level.** Table S5 compares the species-level and regional-level uncertainty of the differences between TabPFN 3.5-FT and the main comparators. The stratified bootstrap resamples species within each region (5,000 resamples). The linear mixed model fits the paired differences with region as a random effect (REML), so its interval reflects variation among regions.

**Table S5.** AUC difference between TabPFN 3.5-FT and each comparator: mean across species, stratified bootstrap 95% interval, mixed-model estimate and 95% interval, and the number of regions and species with a positive difference.

| **Comparator** | **Mean** | **Bootstrap 95%** | **Mixed model** | **Mixed-model 95%** | **Regions** | **Species** |
| --- | --- | --- | --- | --- | --- | --- |
| MaxEnt | +0.0080 | +0.0020 to +0.0138 | +0.0043 | -0.0116 to +0.0203 | 5/6 | 147/225 |
| MaxNet | +0.0075 | +0.0007 to +0.0138 | +0.0034 | -0.0136 to +0.0204 | 5/6 | 140/225 |
| RF down-sampled | +0.0069 | +0.0028 to +0.0114 | +0.0038 | -0.0095 to +0.0172 | 4/6 | 139/225 |
| BRT | +0.0120 | +0.0068 to +0.0176 | +0.0086 | -0.0059 to +0.0230 | 5/6 | 133/225 |
| GAM | +0.0167 | +0.0087 to +0.0247 | +0.0136 | -0.0125 to +0.0396 | 5/6 | 154/225 |
| Ensemble | +0.0038 | -0.0004 to +0.0078 | +0.0012 | -0.0125 to +0.0150 | 5/6 | 115/225 |
| TabPFN 3.5 | +0.0038 | +0.0011 to +0.0067 | +0.0018 | -0.0113 to +0.0149 | 3/6 | 127/225 |

**Calibration.** For each species and model we fitted a logistic regression of observed presence at the survey sites on the logit of the prediction; the slope is the calibration slope (Fig. S9). A slope of 1 indicates that differences in predicted log-odds between sites are correct, and a slope below 1 indicates predictions that are too extreme. The intercept is not interpretable for presence–background models, because they do not estimate prevalence. Averaging over bags shrinks predictions and raises the slope: the median slope was 0.65 and 0.70 for TabPFN 3.5 with 5 and 100 bags, and 0.72 and 0.78 for TabPFN 3.5-FT.

**Variance among species and models.** In a two-way linear model of AUC with species and model as factors (12 single models), species accounted for 89% of the sum of squares and models for 2.6%. With region instead of species, region accounted for 18%.

**Five-bag setting.** With five bags, TabPFN 3.5-FT exceeded MaxEnt by 0.005 (bootstrap 95% interval −0.002 to 0.011; Wilcoxon *P* = 0.04; higher for 130 of 225 species), MaxNet by 0.004 (*P* = 0.14), RF down-sampled by 0.004 (*P* = 0.17) and BRT by 0.009 (*P* = 0.02), and did not differ from the ensemble (*P* = 0.86).

**Encoding of categorical covariates.** With five bags and default weights, TabPFN 3.5 gave a mean AUC of 0.7262 with categorical features and 0.7256 with numeric codes (225 species; the largest regional difference was 0.005 in CAN). The fine-tuning comparisons, which use numeric codes, are therefore not affected by the encoding.

**S6. Spatially separated evaluation**

We followed Valavi et al. (21). Each species was trained on its presence-only records plus the target-group background (TGB) sample of its biological group, using their published samples, and was tested on the independent presence–absence sites. Training records and test sites were assigned to the published spatial blocks by spatial join, and blocks were allocated to 5 folds at random. We kept the most even of 100 allocations, using a port of their spatial_block function (seed 32). An allocation replaced the current best if its smallest count of training presences, training background, test presences or test absences in any fold was at least as large, and the standard deviation of these counts was smaller. Fold *f* was trained on the records outside *f* and tested on the sites inside *f*. Random 5-fold cross-validation drew the same numbers of training presences and background points, and of test presences and absences, at random, as in their code.

A species was excluded if any fold lacked a class, if any training fold had fewer than 5 presences, or if it had fewer than 10 test presences in total. This left 178 species and 1,780 fold tasks. Results are shown in Fig. S14. The covariates were those of Table S1 except NSW vegsys, which Valavi et al. (21) did not use. AUC was computed per fold and averaged per species. The R models used the settings of Section S2. TabPFN used the Fast setting, and TabPFN 3.5-FT used the leave-one-region-out checkpoints, so no checkpoint had seen data from the region it predicted. With target-group background, fine-tuning changed mean AUC in Ontario by +0.003 (random cross-validation) and +0.005 (spatial cross-validation), in contrast to the loss observed with random background in the main benchmark.

**S7. Computation**

All analyses ran on a single NVIDIA DGX Spark (GB10 Grace Blackwell, 20 CPU cores, 128 GB unified memory). The R models (R 4.5.2) were run with 12 species in parallel, one core each, and the foundation models used the GB10 GPU. Fine-tuning one PB model took 2–118 min, and most runs took 8–9 min; run times varied with other jobs that shared the GPU. Because jobs shared the machine, the times in Table S6 are indicative rather than strict benchmarks.

**Table S6.** Run time per species (fit and prediction at all test sites), 225 species.

| **Model** | **Hardware** | **Median (s)** | **Mean (s)** | **Total (h)** |
| --- | --- | --- | --- | --- |
| MaxEnt | CPU, 1 core | 19.5 | 18.3 | 1.1 |
| MaxNet | CPU, 1 core | 24.2 | 66.8 | 4.2 |
| Lasso | CPU, 1 core | 50.8 | 99.5 | 6.2 |
| GAM | CPU, 1 core | 201.6 | 1172.2 | 73.3 |
| BRT | CPU, 1 core | 470.0 | 858.0 | 53.6 |
| RF | CPU, 1 core | 15.7 | 18.1 | 1.1 |
| RF down-sampled | CPU, 1 core | 3.7 | 4.3 | 0.3 |
| XGBoost | CPU, 16 threads | 45.9 | 84.0 | 5.2 |
| Ensemble (sum of members) | CPU | 948.4 | 2152.4 | 134.5 |
| TabICL | GPU | 0.7 | 2.0 | 0.1 |
| TabPFN 2.5, Fast / Stable | GPU | 4.1 / 80.6 | 10.1 / 338.6 | 0.6 / 21.2 |
| TabPFN 3.5, Fast / Stable | GPU | 5.1 / 173.8 | 13.8 / 502.4 | 0.9 / 31.4 |
| TabPFN 3.5-FT, Fast / Stable | GPU | 3.8 / 111.6 | 11.9 / 442.3 | 0.7 / 27.6 |

**S8. Virtual species**

Virtual species were simulated on the 50,000 background points of each region, which carry the real covariate values of the landscape. Ten species were simulated per region with a fixed seed. The niche of each species comprised two or three continuous covariates chosen at random. Each response was unimodal with probability 0.7, a Gaussian function of the standardised covariate with optimum uniform in [−1.5, 1.5] and width log-uniform in [0.4, 1.5], and monotone otherwise, a linear function on the logit scale with slope of absolute value uniform in [1, 3]. The summed linear predictor was rescaled to a standard deviation uniform in [1.5, 3], and the intercept was found by bisection so that the mean probability over the landscape equalled a prevalence log-uniform in [0.03, 0.30]. True occurrence was drawn from a Bernoulli distribution at every point; species with fewer than 300 occupied points outside the test set, or fewer than 20 occupied test points, were redrawn.

Each landscape was split at random into 5,000 test points, 20,000 background points and the remaining points. Presences were drawn without replacement from the occupied points outside the test set. For even-numbered species they were drawn uniformly; for odd-numbered species they were drawn in proportion to (*bz*), where *z* is a standardised covariate outside the niche chosen at random and |*b*| is uniform in [1, 2]. A single sample of 300 presences was drawn and its first 10, 30 and 100 records formed the smaller, nested samples.

For partial dependence, 100 test points were drawn at random, and for each niche covariate these points were predicted with the covariate set in turn to 12 quantiles (2% to 98%) of its landscape distribution. The partial-dependence curve is the mean logit of these predictions, and the true curve is the additive component of the linear predictor at the same quantiles. All models predicted the test points and the partial-dependence points together. The reference models were fitted with the R code used for the benchmark, with the 20,000 background points. TabPFN used 20 balanced bags without test-time augmentation (the first 5 bags give the five-bag results), with default weights and, for TabPFN 3.5-FT, the leave-one-region-out checkpoint of the region in which the species was simulated. Example species in the response-curve figure were chosen, within each category, as the species whose curve recovery by TabPFN 3.5-FT with 30 records was closest to the median of that category.

**Table S7.** Virtual species: mean Spearman correlation with the true probability, AUC against simulated presences and absences, and response-curve recovery, for 10 / 30 / 100 / 300 presence records (60 species per design).

| **Model** | **Spearman with truth** | **AUC** | **Curve recovery** |
| --- | --- | --- | --- |
| *Additive relationships (AUC of the true probability 0.79)* | | | |
| TabPFN 3.5-FT | 0.48 / 0.57 / 0.64 / 0.71 | 0.68 / 0.70 / 0.71 / 0.73 | 0.51 / 0.64 / 0.74 / 0.79 |
| TabPFN 3.5 | 0.47 / 0.58 / 0.66 / 0.73 | 0.67 / 0.70 / 0.72 / 0.73 | 0.44 / 0.59 / 0.71 / 0.79 |
| MaxEnt | 0.51 / 0.64 / 0.68 / 0.75 | 0.68 / 0.71 / 0.72 / 0.74 | 0.54 / 0.72 / 0.80 / 0.88 |
| MaxNet | 0.48 / 0.63 / 0.69 / 0.75 | 0.66 / 0.70 / 0.72 / 0.74 | 0.54 / 0.73 / 0.81 / 0.91 |
| GAM | 0.46 / 0.61 / 0.70 / 0.76 | 0.65 / 0.70 / 0.72 / 0.74 | 0.51 / 0.64 / 0.84 / 0.91 |
| Lasso | 0.46 / 0.63 / 0.72 / 0.76 | 0.67 / 0.71 / 0.73 / 0.74 | 0.41 / 0.61 / 0.84 / 0.92 |
| BRT | 0.43 / 0.58 / 0.64 / 0.72 | 0.67 / 0.70 / 0.71 / 0.73 | 0.35 / 0.58 / 0.77 / 0.84 |
| RF down-sampled | 0.49 / 0.56 / 0.60 / 0.66 | 0.68 / 0.69 / 0.71 / 0.72 | 0.48 / 0.55 / 0.63 / 0.72 |
| RF | 0.12 / 0.21 / 0.34 / 0.48 | 0.53 / 0.57 / 0.63 / 0.67 | 0.18 / 0.19 / 0.29 / 0.37 |
| Ensemble | 0.54 / 0.64 / 0.70 / 0.75 | 0.69 / 0.71 / 0.73 / 0.74 | 0.62 / 0.71 / 0.85 / 0.90 |
| *Complex relationships (AUC of the true probability 0.80)* | | | |
| TabPFN 3.5-FT | 0.40 / 0.48 / 0.59 / 0.69 | 0.65 / 0.67 / 0.70 / 0.72 | 0.43 / 0.51 / 0.66 / 0.69 |
| TabPFN 3.5 | 0.39 / 0.49 / 0.64 / 0.74 | 0.65 / 0.67 / 0.71 / 0.73 | 0.39 / 0.46 / 0.64 / 0.69 |
| MaxEnt | 0.41 / 0.53 / 0.64 / 0.71 | 0.63 / 0.68 / 0.71 / 0.73 | 0.37 / 0.59 / 0.73 / 0.75 |
| MaxNet | 0.39 / 0.52 / 0.66 / 0.73 | 0.63 / 0.67 / 0.71 / 0.73 | 0.36 / 0.60 / 0.73 / 0.75 |
| GAM | 0.42 / 0.53 / 0.61 / 0.68 | 0.63 / 0.67 / 0.70 / 0.72 | 0.42 / 0.49 / 0.70 / 0.68 |
| Lasso | 0.39 / 0.49 / 0.61 / 0.67 | 0.63 / 0.67 / 0.70 / 0.72 | 0.34 / 0.48 / 0.67 / 0.74 |
| BRT | 0.32 / 0.47 / 0.60 / 0.70 | 0.63 / 0.67 / 0.71 / 0.73 | 0.28 / 0.49 / 0.64 / 0.67 |
| RF down-sampled | 0.41 / 0.48 / 0.56 / 0.64 | 0.65 / 0.68 / 0.70 / 0.72 | 0.39 / 0.47 / 0.56 / 0.63 |
| RF | 0.12 / 0.20 / 0.34 / 0.47 | 0.53 / 0.57 / 0.63 / 0.67 | 0.15 / 0.23 / 0.30 / 0.40 |
| Ensemble | 0.44 / 0.52 / 0.63 / 0.71 | 0.65 / 0.68 / 0.71 / 0.73 | 0.46 / 0.56 / 0.70 / 0.71 |

**Complex relationships.** A second set of 60 species (ten per region, seed as above) was simulated after the first set had been analysed, to test whether the additive design favoured regression-based methods. Each species responded jointly to two continuous covariates in one of three ways, chosen with equal probability: a bivariate Gaussian with correlated tolerances (centres uniform in [−1.2, 1.2], tolerances log-uniform in [0.5, 1.5], correlation of absolute value uniform in [0.5, 0.85]; 24 species); a joint threshold, 4 (6 *d*_1_ (*z*_1_ − *t*_1_)) (6 *d*_2_ (*z*_2_ − *t*_2_)) with thresholds uniform in [−1, 1] and random directions *d* (16 species); or a multiplicative interaction, β *z*_1_ *z*_2_ + γ *z*_1_ with |β| uniform in [1, 2] and γ uniform in [−1, 1] (20 species). With probability 0.5, a third covariate had an additive Gaussian effect. Scaling, prevalence, sampling bias, nested samples and models were as for the additive design. True partial responses were computed by applying the true function to the same partial-dependence points used for the models.

**Interpolation and extrapolation.** For each dataset, test points were classified as inside the environmental envelope of the presences if they lay within the range of the presence records on every niche covariate. The fraction of test points inside the envelope was 0.40, 0.62, 0.78 and 0.85 with 10, 30, 100 and 300 records. Spearman correlations recomputed inside the envelope (231 datasets with more than 50 such points; additive design) are reported in the main text.

**Bias and covariate correlation.** For the 30 additive species with environmental sampling bias, we computed the largest absolute correlation over the landscape between the bias covariate and any niche covariate (median 0.48, range 0.07–0.84), and regressed the difference between TabPFN 3.5-FT and MaxEnt on this correlation and on the logarithm of the number of records. The slope was not significant for response-curve recovery (*P* = 0.24) or for the Spearman correlation (*P* = 0.84).

**Calibration against simulated presences and absences.** The median calibration slope (additive design, all sample sizes) was 0.96 for TabPFN 3.5-FT, 0.98 for TabPFN 3.5, 1.05 for BRT, 0.86 for Lasso, 0.71 for RF down-sampled, 0.66 for the ensemble, 0.43 for MaxEnt, 0.38 for MaxNet and 0.30 for GAM. For TabPFN 3.5-FT it was 1.24, 0.98, 0.90 and 0.82 with 10, 30, 100 and 300 records, and for MaxEnt 0.37, 0.35, 0.45 and 0.58.

**Sensitivity to the number of background points per bag.** Because each balanced bag contains only as many background points as presences, the foundation model sees little of the landscape when records are few. To test whether this explains its weaker niche recovery, we refitted TabPFN 3.5-FT to the datasets with 10 and 30 presences with five times as many background points as presences in each bag (20 bags). This made all three measures worse. With 10 and 30 records, the Spearman correlation with the true probability changed by −0.038 (*P* = 0.002) and −0.008 (*P* = 0.14), AUC by −0.008 (*P* = 0.008) and −0.002 (*P* = 0.62), and response-curve recovery by −0.22 and −0.20 (both *P* < 10^−4^; paired Wilcoxon tests across 60 species). Scarcity of background points in the bags therefore does not explain the deficit, and the balanced design remains the better choice, as on the benchmark (Table S2).

**S9. Supplementary figures**

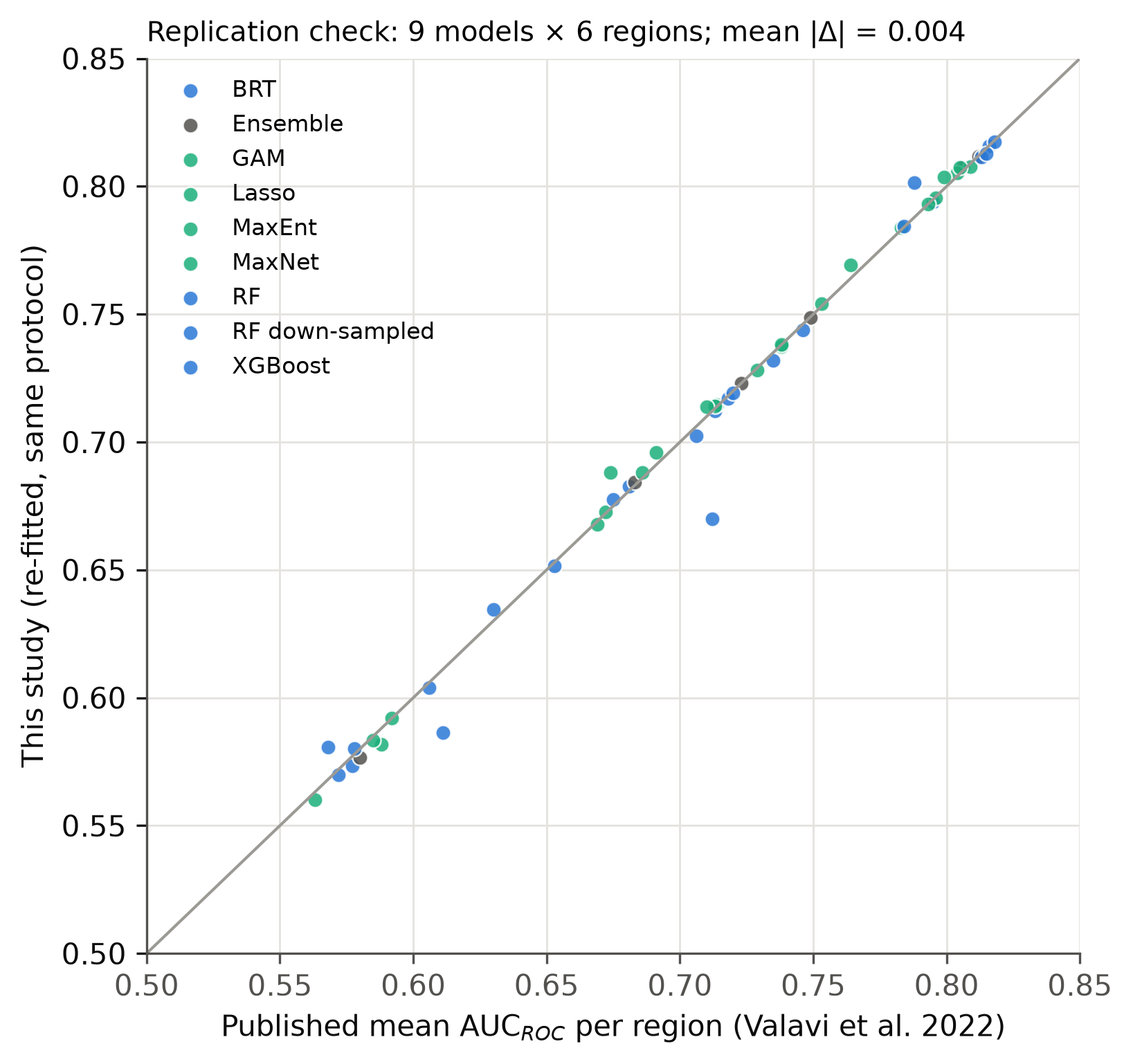

**Fig. S1.** Replication check. Regional mean AUC of our re-fitted reference models against the values published by Valavi et al. (3).

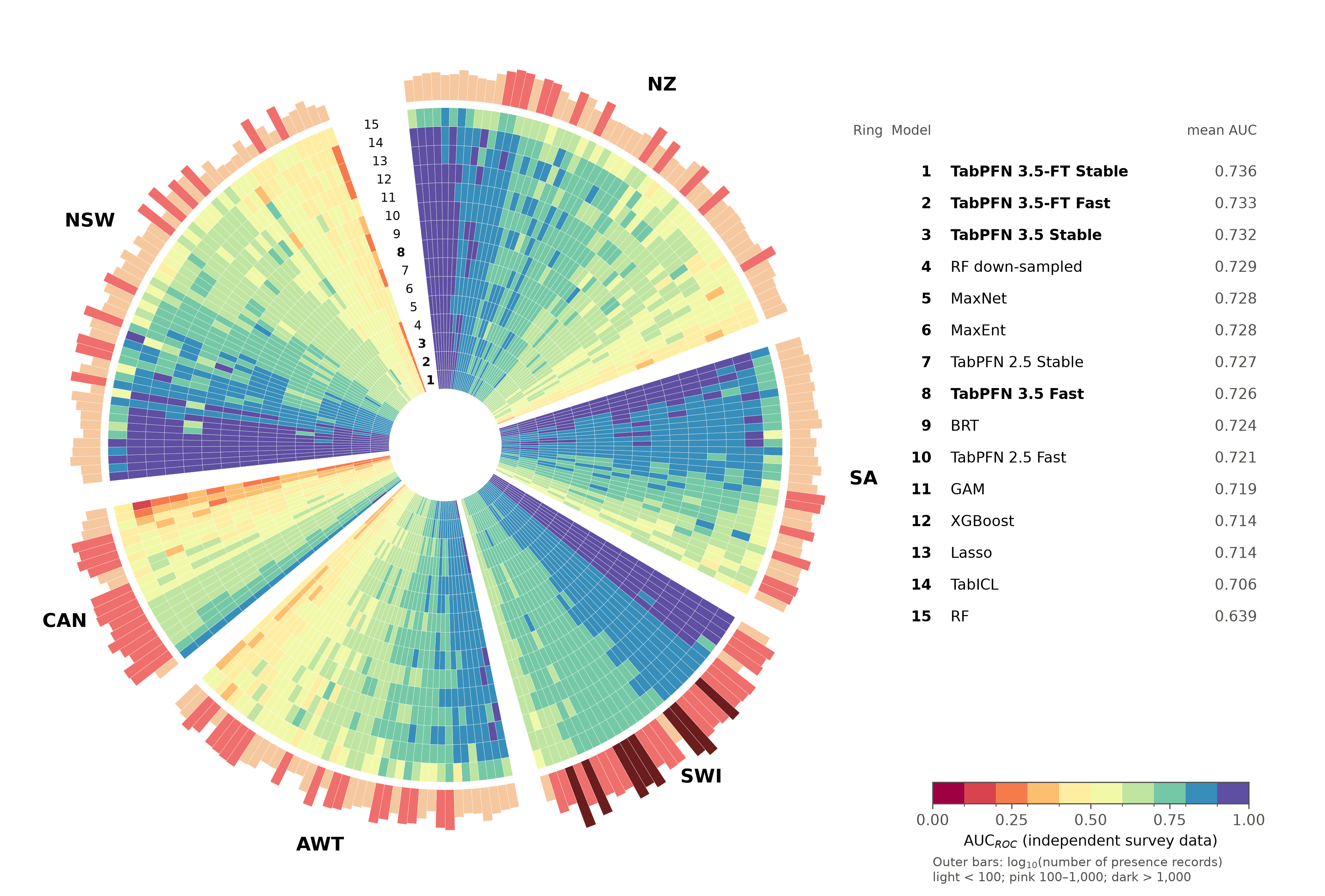

**Fig. S2.** Per-species AUC of all models and variants except ensembles, including the five-bag (Fast) TabPFN settings. Layout as in Fig. 1 of the main text: ring numbers in the gap follow the key, which lists models by mean AUC.

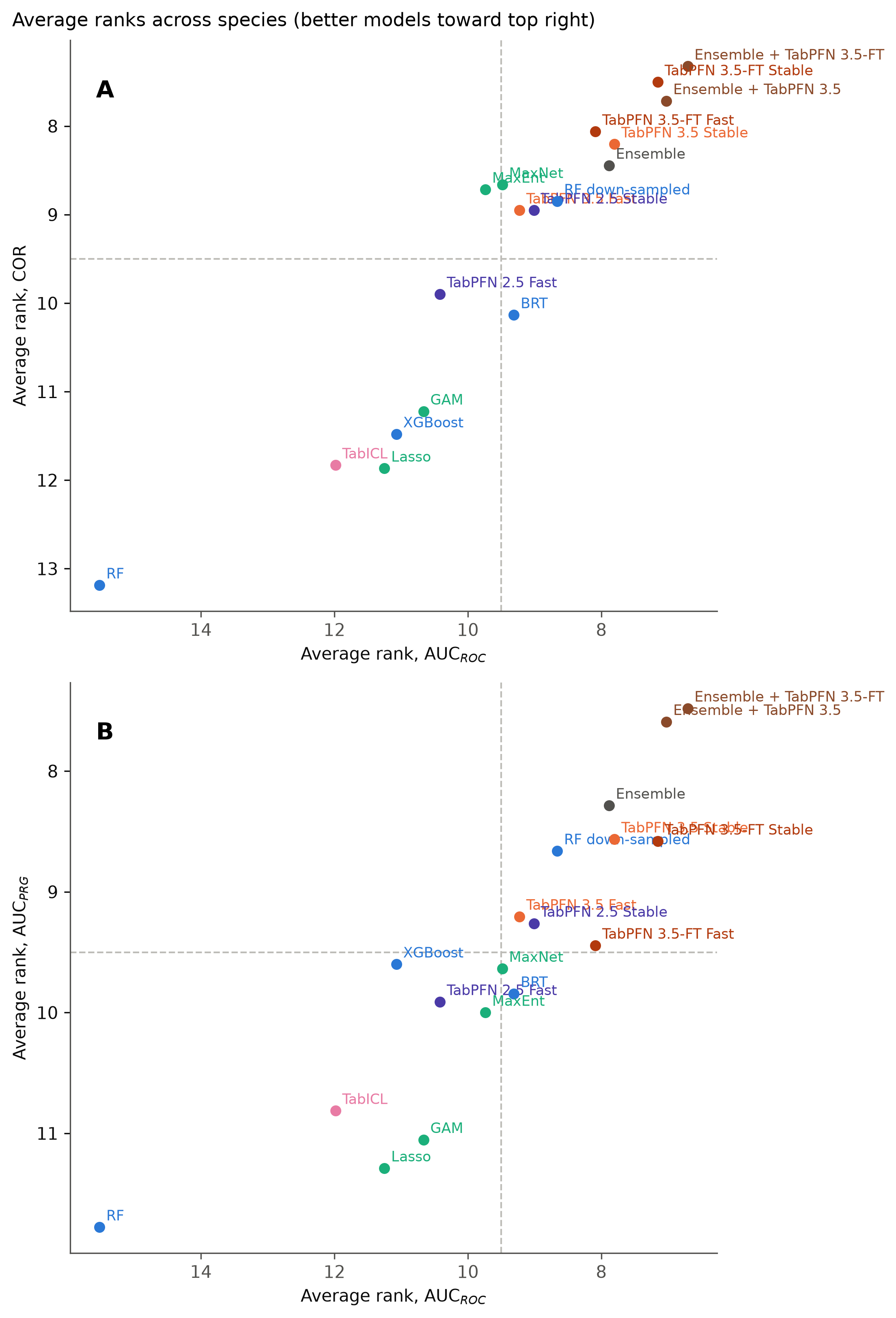

**Fig. S3.** Average ranks across species for all models and variants, including ensembles that contain TabPFN. (A) AUC against COR. (B) AUC against AUC_PRG_. Better models lie towards the top right.

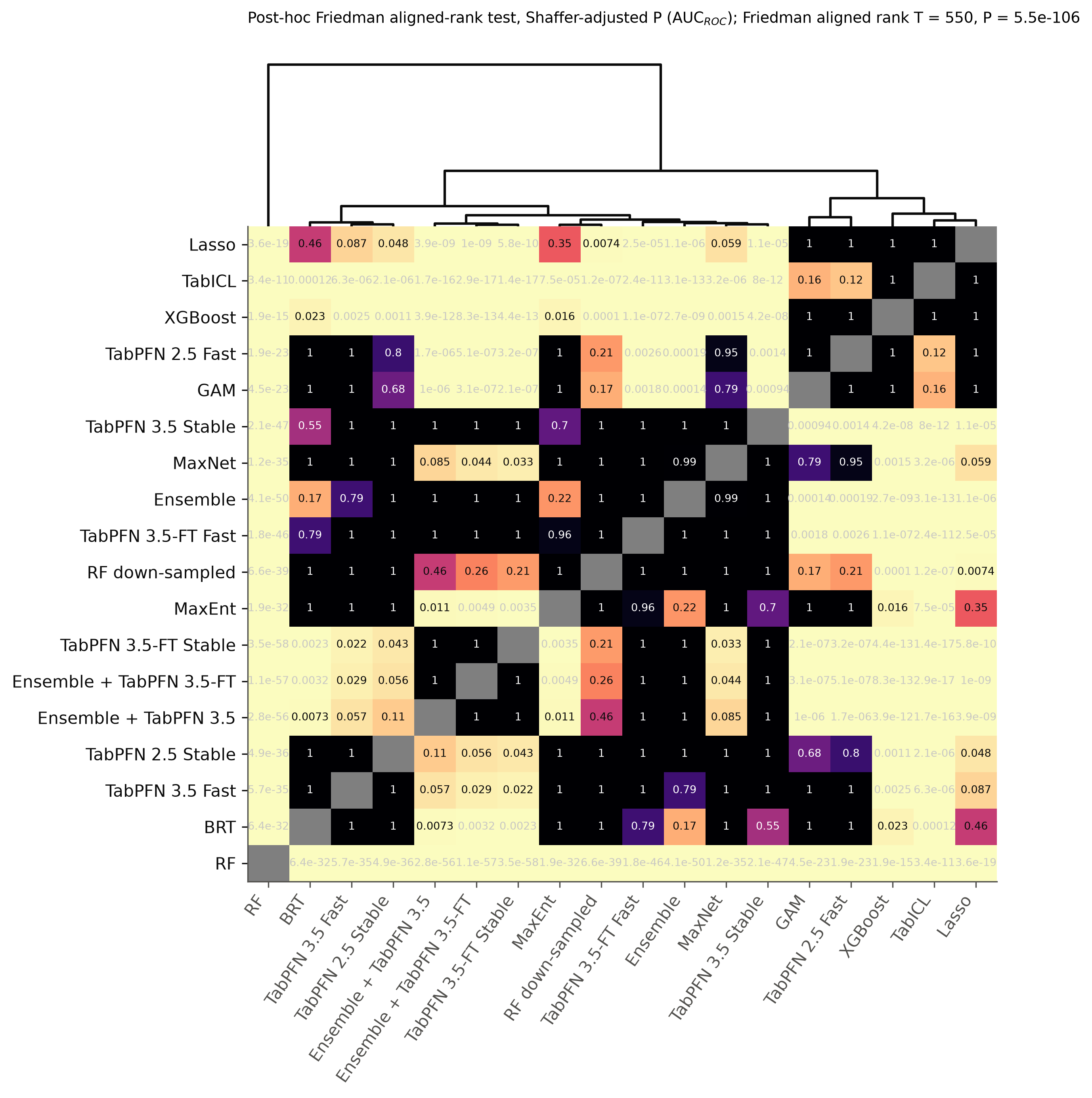

**Fig. S4.** Shaffer-corrected *P* values of pairwise aligned-rank post hoc tests on AUC, for all models and variants (18 approaches).

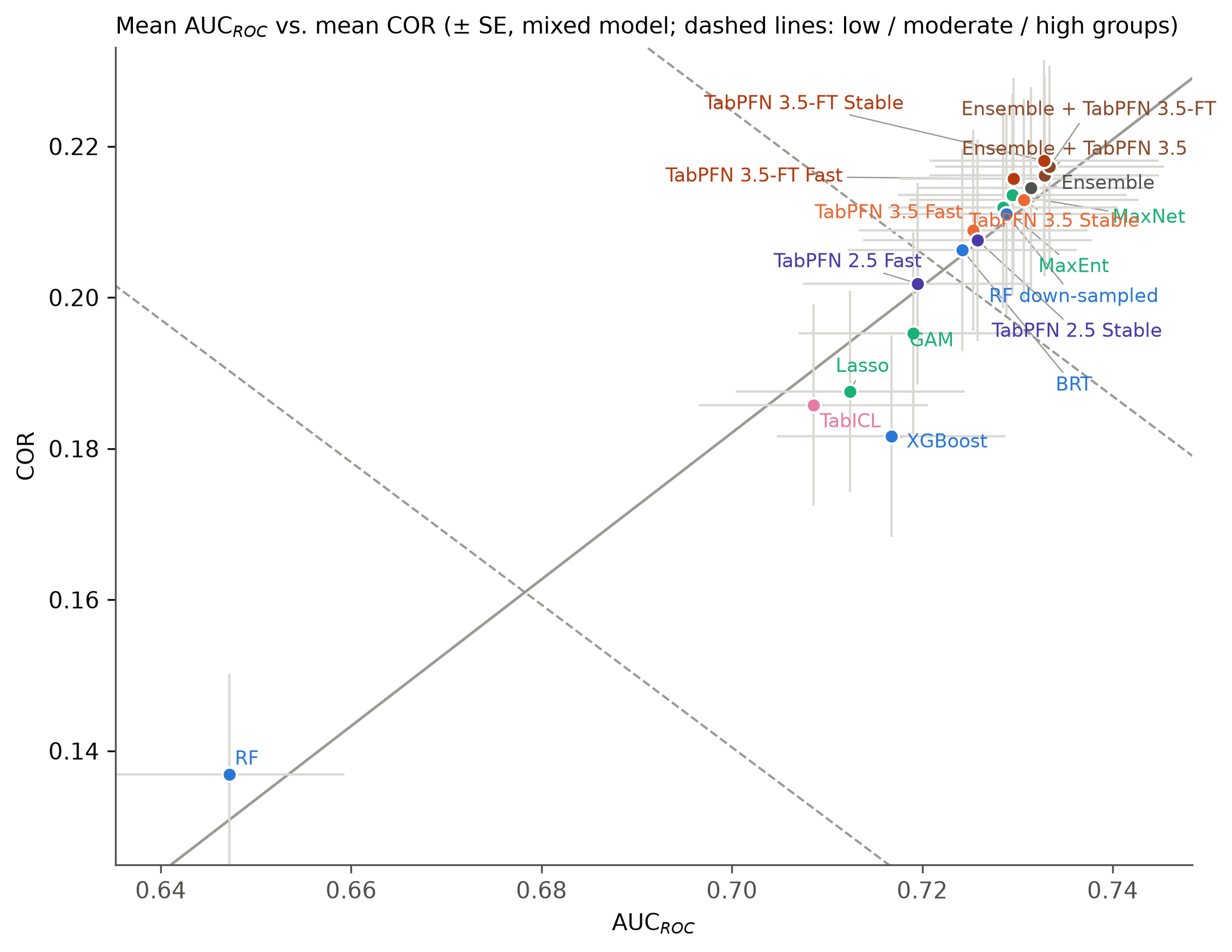

**Fig. S5.** Mean AUC against mean COR, with performance groups from complete-linkage clustering, as in Valavi et al. (3).

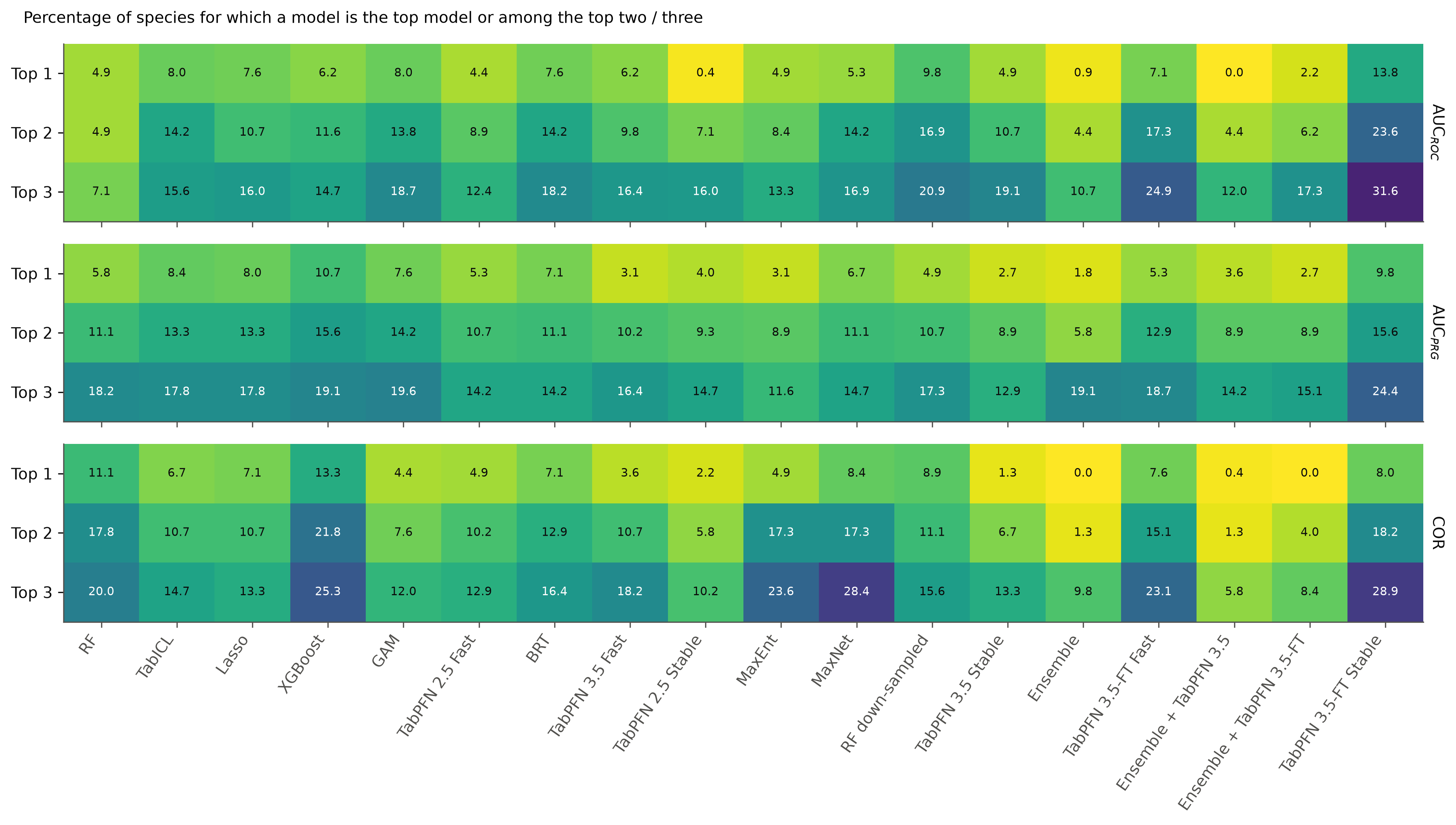

**Fig. S6.** Percentage of species for which each model is the best, among the top two, or among the top three, for AUC, AUC_PRG_ and COR.

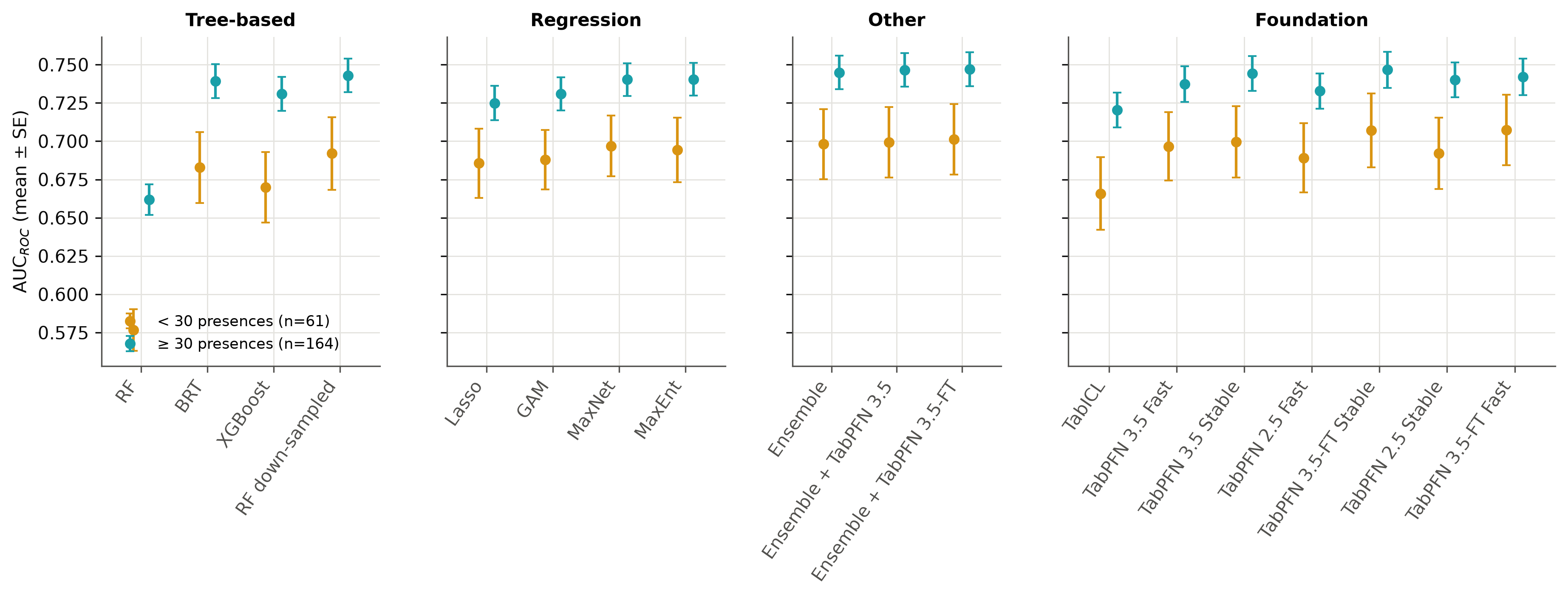

**Fig. S7.** Mean AUC (± SE) for species with fewer than 30 (*n* = 61) and with 30 or more (*n* = 164) presence records, all models.

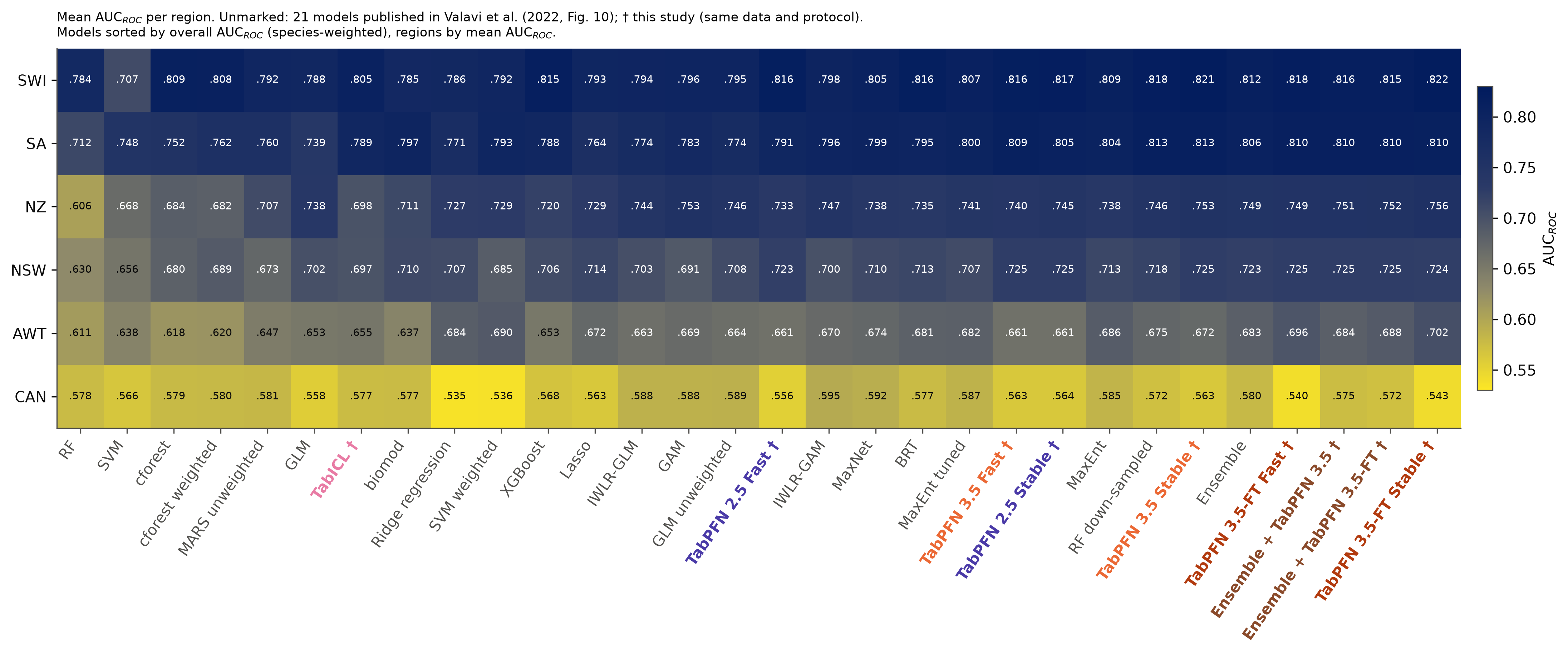

**Fig. S8.** Regional mean AUC of the 21 published methods (3) and all TabPFN variants and ensembles (†).

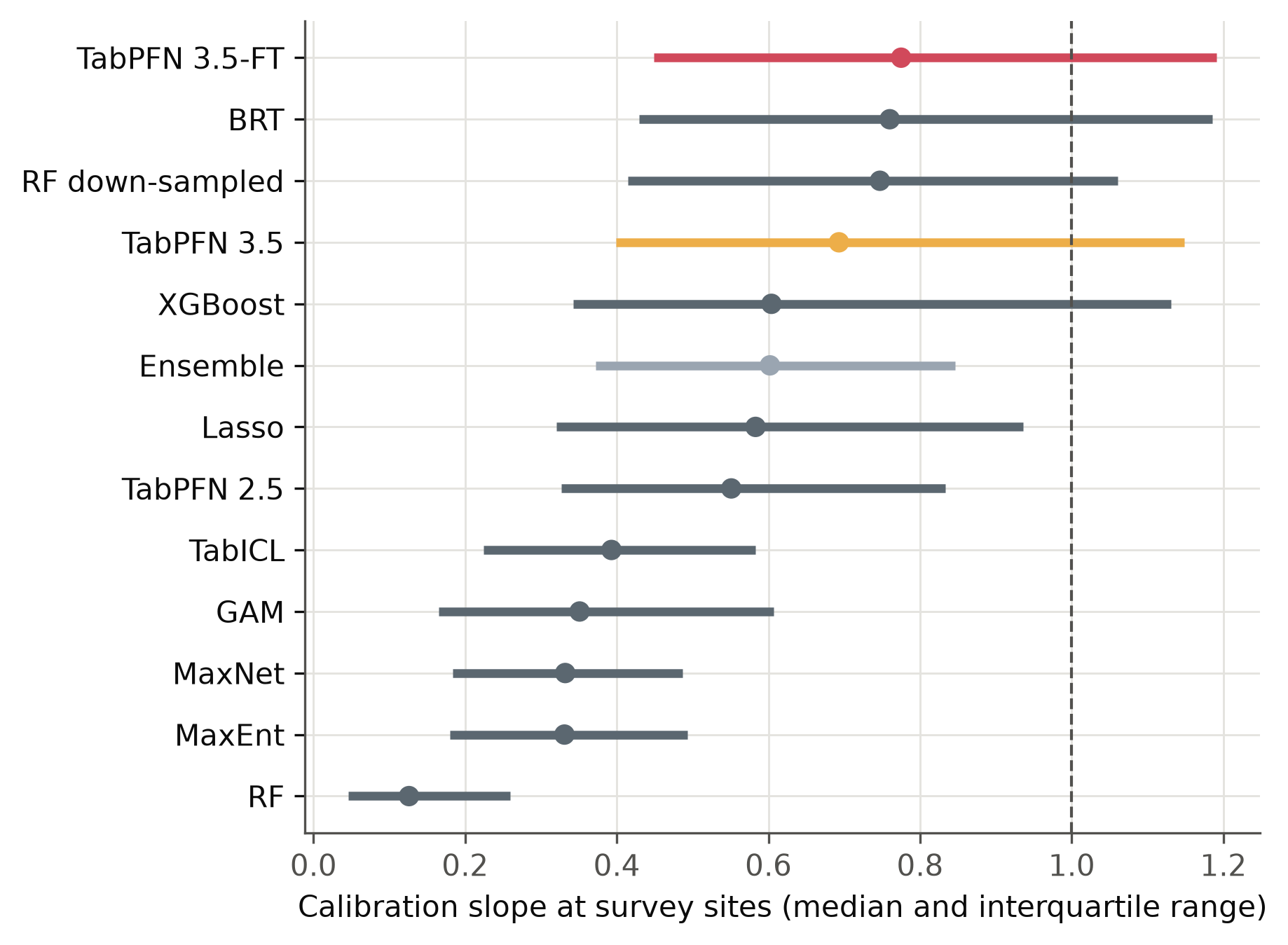

**Fig. S9.** Calibration slope at the independent survey sites (median and interquartile range across species). The dashed line marks perfect calibration of relative likelihood.

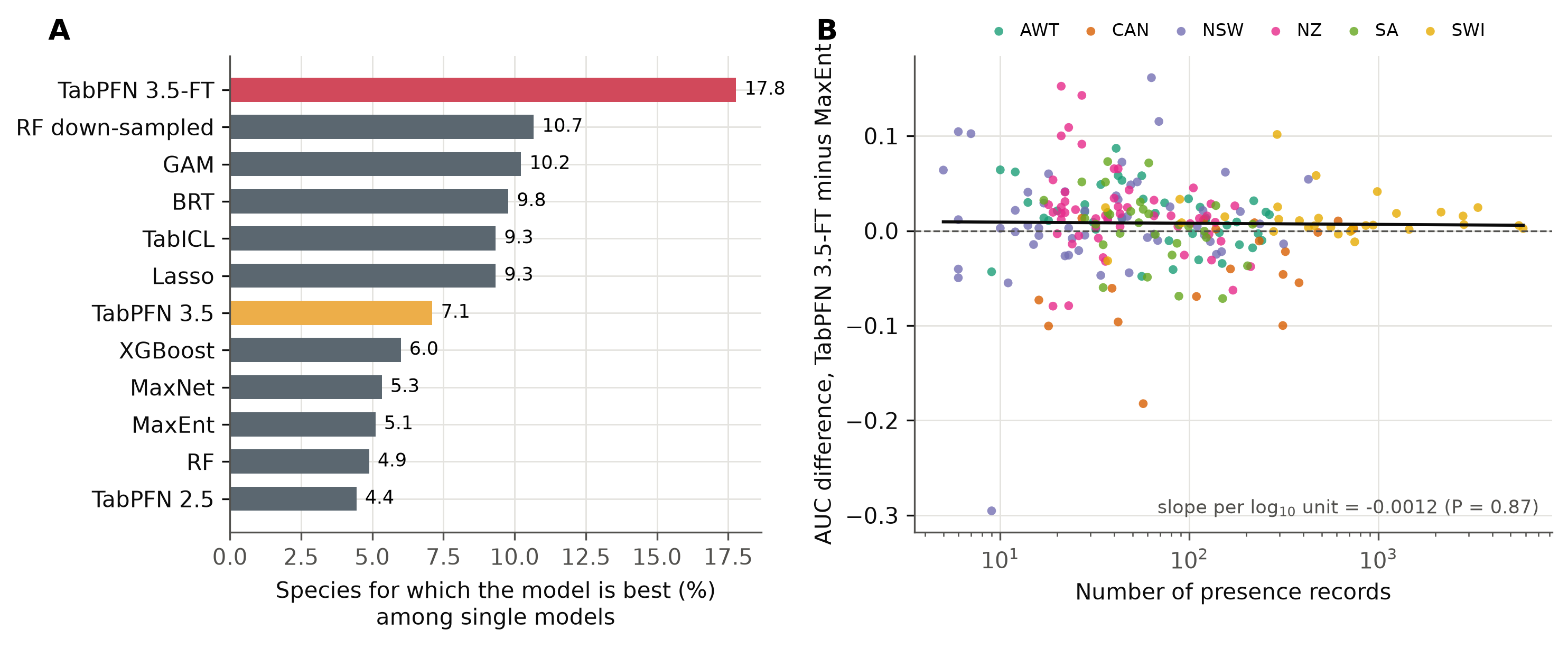

**Fig. S10.** Species-level results. (A) Share of species for which each single model had the highest AUC; ties are shared. (B) Difference in AUC between TabPFN 3.5-FT and MaxEnt against the number of presence records (log scale), coloured by region. The line shows the slope from a regression with region as a fixed effect, which was not significant.

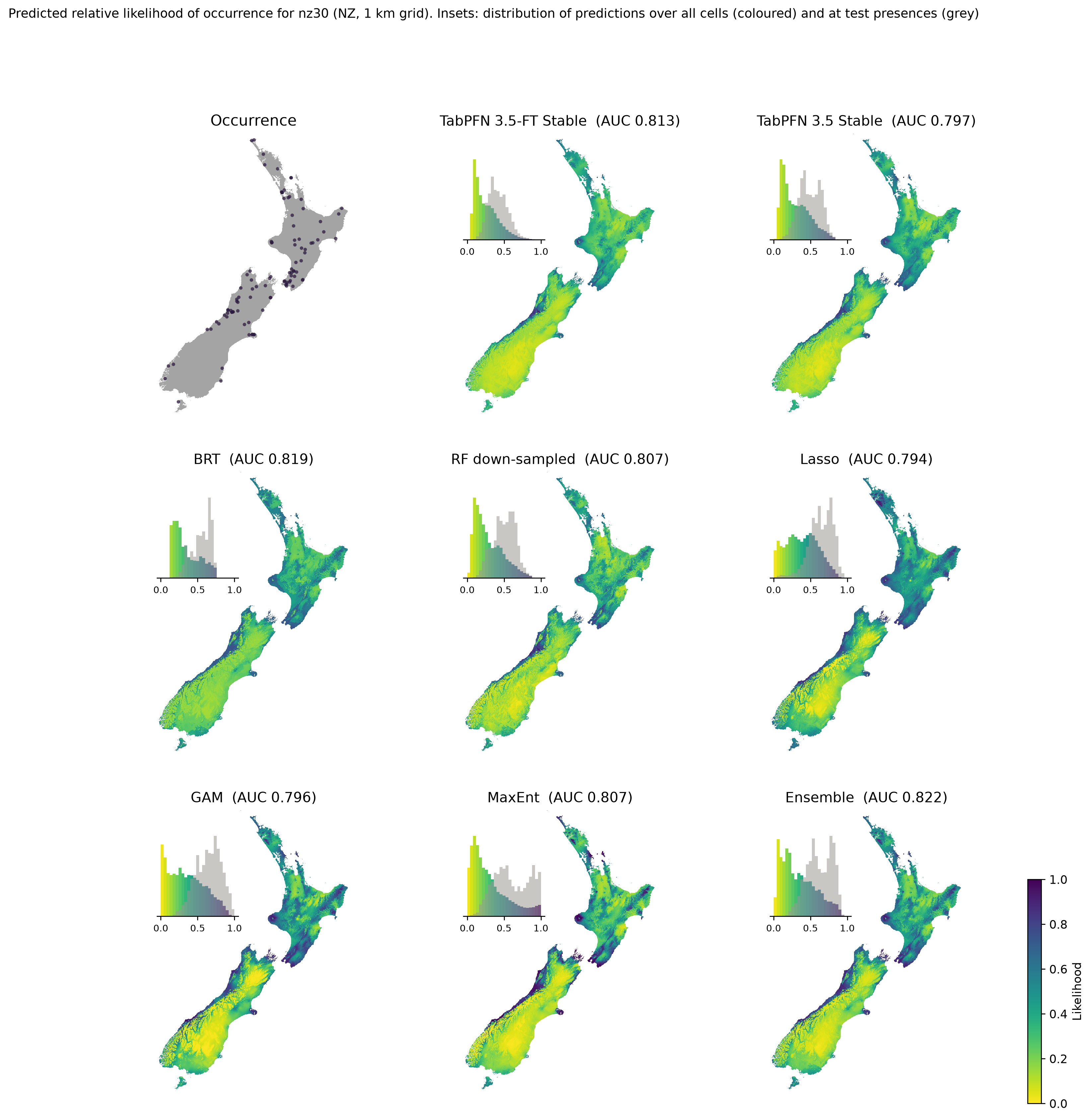

**Fig. S11.** Predicted relative likelihood of occurrence of the New Zealand species nz30, the example used by Valavi et al. (3), on a 1-km grid from eight models, with AUC on the independent survey data. The top-left panel shows the presence records. Insets show the distribution of predictions over all cells (coloured) and at test presences (grey).

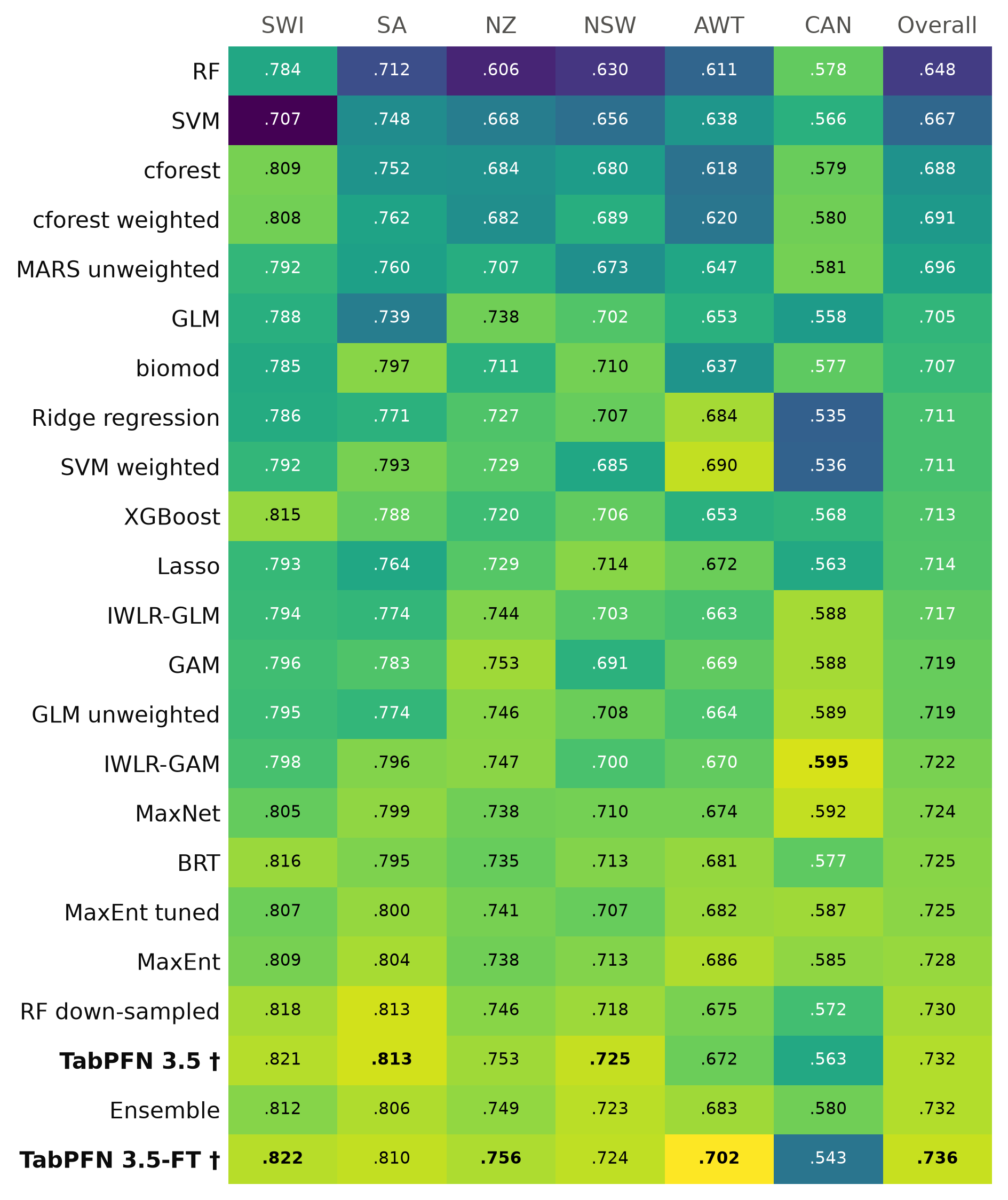

**Fig. S12.** Regional mean AUC of TabPFN and the 21 published methods. Values for the 21 methods are from Valavi et al. (3). † marks TabPFN 3.5 with and without fine-tuning (100 bags). Rows are sorted by the species-weighted overall mean. Cells are coloured by the z-score within each column, and the best value in each column is in bold.

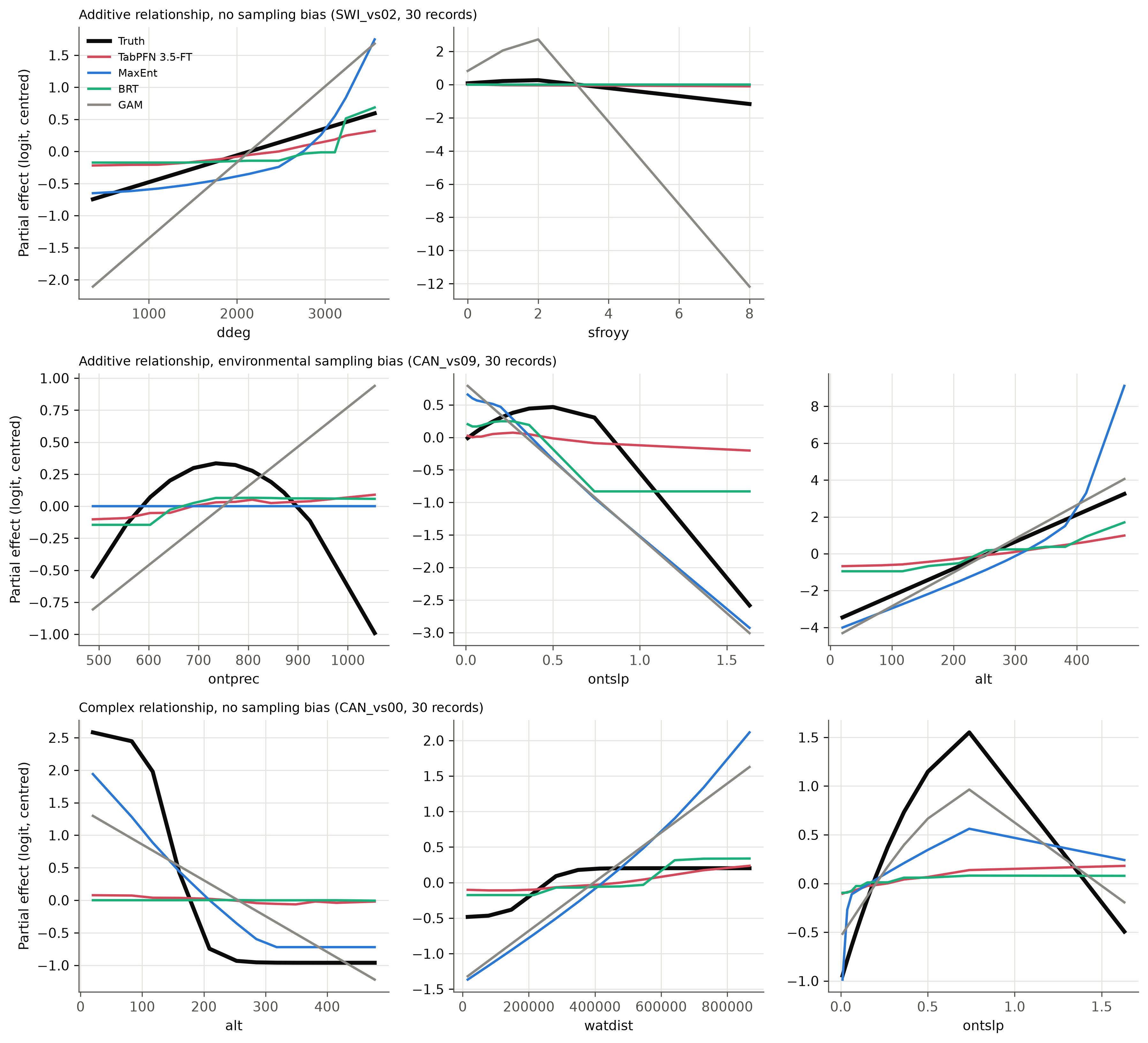

**Fig. S13.** Examples of true and estimated partial responses (centred, logit scale) for virtual species fitted to 30 presence records: an additive species without sampling bias (top), an additive species with environmental sampling bias (middle) and a complex species without bias (bottom). Each example is the species of its category whose response-curve recovery by TabPFN 3.5-FT was closest to the category median.

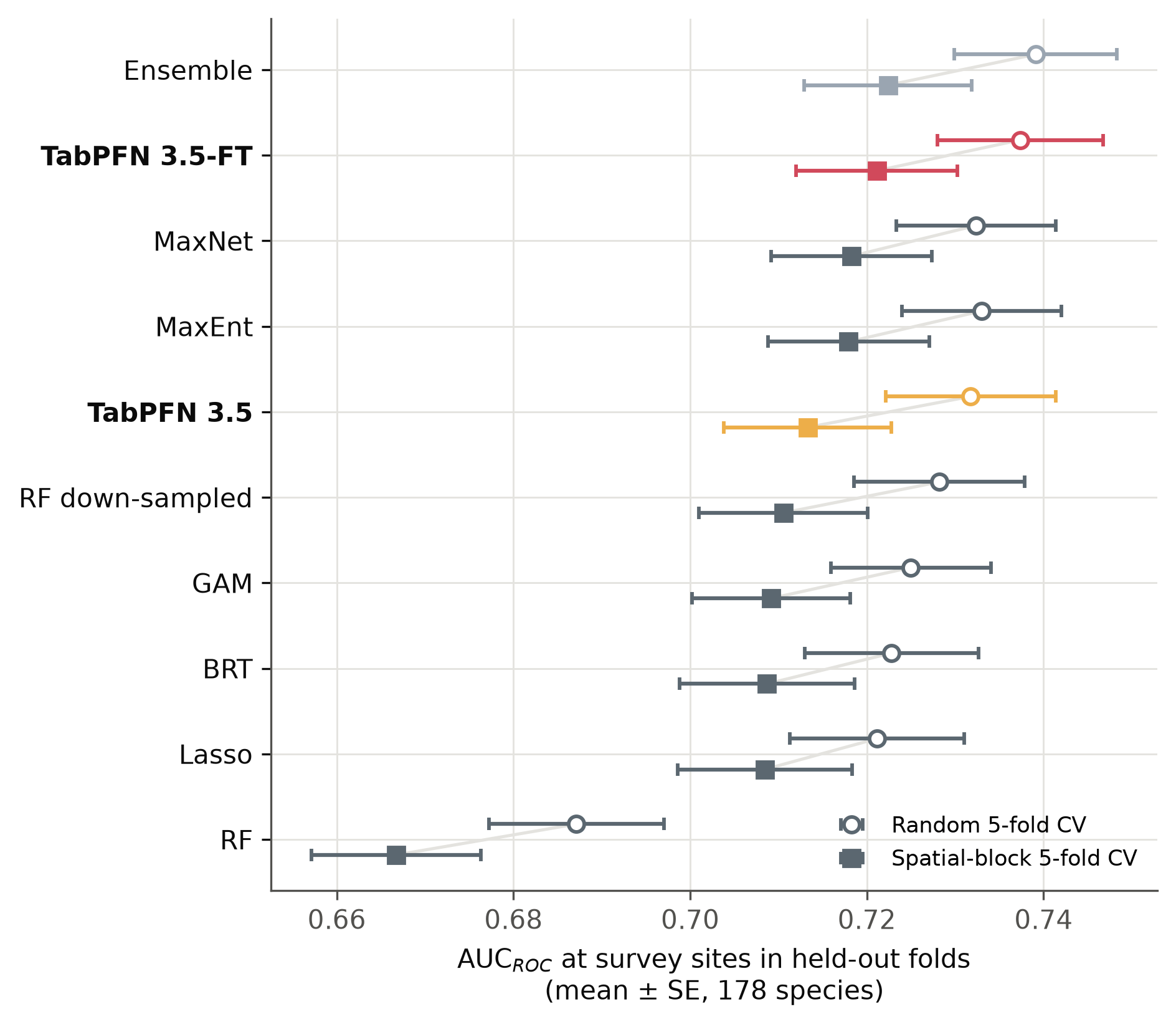

**Fig. S14.** Spatially separated evaluation. Mean AUC (± SE, 178 species) under random (circles, upper) and spatial-block (squares, lower) 5-fold cross-validation (21). Models were trained on presence records with target-group background and tested on the independent survey sites in held-out folds. Lines connect the two designs for each model. TabPFN models used five bags (Fast setting).
